# Sensitize drug-resistant tumors to chemo- and targeted therapies by smart ferroptosis

**DOI:** 10.64898/2026.11.17.767181

**Authors:** Yunqi Zhao, Lu Wang, Ziyan Kong, Aiwei Yan, Xuhui Wen, Xingyi Xie, Zhimin Fan, Yongqian Shu, Meng Cao, Jinke Wang

**Author notes:** To whom correspondence should be addressed: Meng Cao, Division of Breast Surgery, Department of General Surgery, Nanjing Drum Tower Hospital, the Affiliated Hospital of Medical School, Nanjing University, Nanjing, 210008, China, Jinke Wang, Sipailou 2, School of Biological Science and Medical Engineering, Southeast University, Nanjing 210096, China.

## Abstract

Therapeutic resistance and insufficient sensitivity of tumors to anticancer agents remain major challenges in the current cancer treatments. Here, we developed an NF-κB-responsive rAAV-T7 platform that specifically disrupts ferroptosis and redox defenses in tumor cells by simultaneously targeting seven regulators of ferroptosis defense, redox homeostasis, and iron metabolism, including GPX4, SLC7A11, FSP1, FPN, NRF2, LCN2, and FTH1. rAAV-T7 specifically induced expression of the encoded miRNAs in tumor cells with higher RELA expression, leading to suppression of their corresponding target genes, while showing little or no detectable activity in normal cells. rAAV-T7 enhanced tumor-cell sensitivity to multiple chemotherapeutic agents and restored chemosensitivity in doxorubicin- and gemcitabine-resistant cells. Combination treatment increased intracellular Fe²⁺ accumulation, lipid peroxidation, and total ROS levels, whereas ferrostatin-1 and N-acetylcysteine attenuated the associated cytotoxicity, indicating that ferroptosis-associated oxidative stress contributes to the enhanced therapeutic response. The therapeutic effects of rAAV-T7 were further validated in MCF-7/ADR tumor spheroids and patient-derived colorectal cancer organoids. In vivo, rAAV-T7 potentiated doxorubicin-mediated tumor suppression and restored DOX sensitivity without detectable additional toxicity in major organs under the conditions tested. Moreover, rAAV-T7 sensitized sorafenib-resistant PLC-PRF-5/SR cells to sorafenib, accompanied by enhanced Fe²⁺ accumulation, lipid peroxidation, and oxidative stress. Collectively, these findings establish tumor-selective disruption of ferroptosis and redox defenses as a strategy for enhancing anticancer therapy and overcoming acquired drug resistance.

## Introduction

Advances in chemotherapy, molecular targeted therapy, immunotherapy, and their rational combinations have substantially improved cancer treatment. However, clinical efficacy remains limited by inadequate treatment responses and the development of therapeutic resistance. Some tumors exhibit intrinsically low sensitivity to anticancer agents owing to genetic and epigenetic heterogeneity, altered drug transport and metabolism, impaired cell-death responses, and metabolic adaptation^1–3^. In tumors that initially respond to therapy, adaptive cellular responses and clonal selection can further drive acquired resistance, leading to disease progression and treatment failure^1,2^. Dysregulation of DNA repair, redox homeostasis, and stress-response pathways can further impair treatment responses and promote acquired resistance^2–5^. Although next-generation agents and rational combinations can overcome specific resistance mechanisms, their durable efficacy may be compromised by the activation of bypass signaling pathways and the emergence of additional resistance mechanisms^6,7^. Thus, an unresolved challenge is to enhance tumor susceptibility to treatment despite the diversity and adaptability of resistance mechanisms.

Ferroptosis is an iron-dependent form of regulated cell death characterized by the accumulation of lipid peroxides, distinguishing it mechanistically from apoptosis and other canonical forms of cell death^8,9^. Increasing evidence suggests that ferroptosis has considerable potential to overcome acquired therapeutic resistance. For example, targeting ferroptosis-regulatory pathways has been shown to restore tamoxifen sensitivity in estrogen receptor-positive breast cancer and gemcitabine sensitivity in pancreatic cancer models^10,11^. Tumor cells resist ferroptosis through multiple interconnected defense mechanisms, including the SLC7A11–glutathione–GPX4 axis, the FSP1–CoQ10 system, NRF2-dependent antioxidant programs, and other pathways regulating iron homeostasis and lipid metabolism^8,9,12,13^. Disruption of these defense mechanisms can promote lipid peroxide accumulation and thereby increase the susceptibility of tumor cells to ferroptosis^14,15^. These findings have raised considerable interest in targeting ferroptosis defenses to enhance anticancer therapy. However, several challenges remain in translating ferroptosis-based strategies into effective and selective cancer therapies.

First, insufficient tumor selectivity may increase the risk of ferroptosis-associated toxicity in normal tissues. Ferroptosis is governed by fundamental cellular processes, including iron metabolism, glutathione homeostasis, antioxidant defense, and lipid metabolism, which are also essential for the survival and function of normal tissues^12,15,16^. Second, tumor sensitivity to ferroptosis is highly heterogeneous. Differences in intracellular iron availability, glutathione metabolism, GPX4 and FSP1 activity, lipid composition, and the expression or activity of enzymes involved in polyunsaturated fatty acid metabolism can substantially influence lipid peroxide accumulation and determine the ferroptotic threshold of tumor cells^8,9,12,13,17^. Consequently, ferroptosis-targeting strategies may exhibit variable efficacy across tumor types and even among patients within the same tumor. Importantly, tumor cells possess multiple interconnected ferroptosis-defense pathways that can compensate for the disruption of individual regulatory nodes^8,9,12,13,15^. Thus, targeting a single ferroptosis-defense pathway may be insufficient to overcome the compensatory mechanisms that sustain ferroptosis resistance. Collectively, these limitations highlight the need for strategies that combine tumor-selective activity with simultaneous disruption of multiple ferroptosis-defense mechanisms. Such an approach may help address the heterogeneous and compensatory nature of ferroptosis defense while limiting interference with ferroptosis-protective processes in normal tissues.

To address these challenges, we developed an NF-κB-responsive rAAV-T7 platform for tumor-selective disruption of ferroptosis defenses. NF-κB-responsive regulatory systems have been used to achieve cancer cell-selective therapeutic gene expression, and previous studies have demonstrated the feasibility of decoy and minimal promoter (DMP)-based strategies for ferroptosis gene interference and tumor immunotherapy^18–21^. Building on this principle, rAAV-T7 simultaneously targets seven key regulators of ferroptosis, redox homeostasis, and iron metabolism, including GPX4, SLC7A11, FSP1, FPN, NRF2, LCN2, and FTH1, thereby disrupting multiple ferroptosis-defense pathways. We evaluated rAAV-T7 across multiple tumor models, including three-dimensional tumor spheroids, patient-derived cancer organoids, and xenografts, to assess its therapeutic efficacy, ferroptosis-associated effects, tumor selectivity, biodistribution, and safety. Collectively, our findings establish a tumor-selective ferroptosis that can enhance anticancer efficacy of commonly used chemotherapy and targeted therapy drugs. The smart ferroptosis provides a new tool for overcoming resistance of cancers to those first-line drugs.

## Results

### Elevated RELA expression is associated with tumor-selective expression of pAAV-T7

To determine whether the differential NF-κB signaling context between tumor and normal cells could be exploited to achieve tumor-selective transgene expression, we first constructed the NF-κB-responsive pAAV-T7 system, in which expression of the encoded miRNAs is driven by an NF-κB-responsive regulatory element (Fig. 1a). We first examined RELA/p65 mRNA expression in representative tumor and normal cell lines. qPCR analysis revealed significantly higher RELA expression in tumor cell lines, including MCF-7/ADR, AsPC-1/GEM, PLC-PRF-5/SR, HCT116, MDA-MB-231, and SKOV3, than in normal cell lines, including MCF-12A, MRC-5, HL7702, and GES-1 (Fig. 1b). We next investigated whether this differential RELA expression was associated with selective expression of the encoded miRNAs. Tumor and normal cells were transfected with pAAV-T7 or the empty control vector pAAV-NT, followed by qPCR analysis of the seven encoded miRNAs and their corresponding target genes. Compared with pAAV-NT, pAAV-T7 induced robust expression of all seven miRNAs in tumor cells, accompanied by significant downregulation of their corresponding target genes (Fig. 1c, d, Fig. S1a, b). In contrast, pAAV-T7 did not induce detectable expression of the seven miRNAs in normal cells (Fig. 1e, Fig. S1c). Together, these findings demonstrate that pAAV-T7 specifically drives expression of the encoded miRNAs and reduces the expression of their target genes in tumor cells, supporting the tumor-selective activity of its NF-κB-responsive regulatory element.

**Fig.1:**
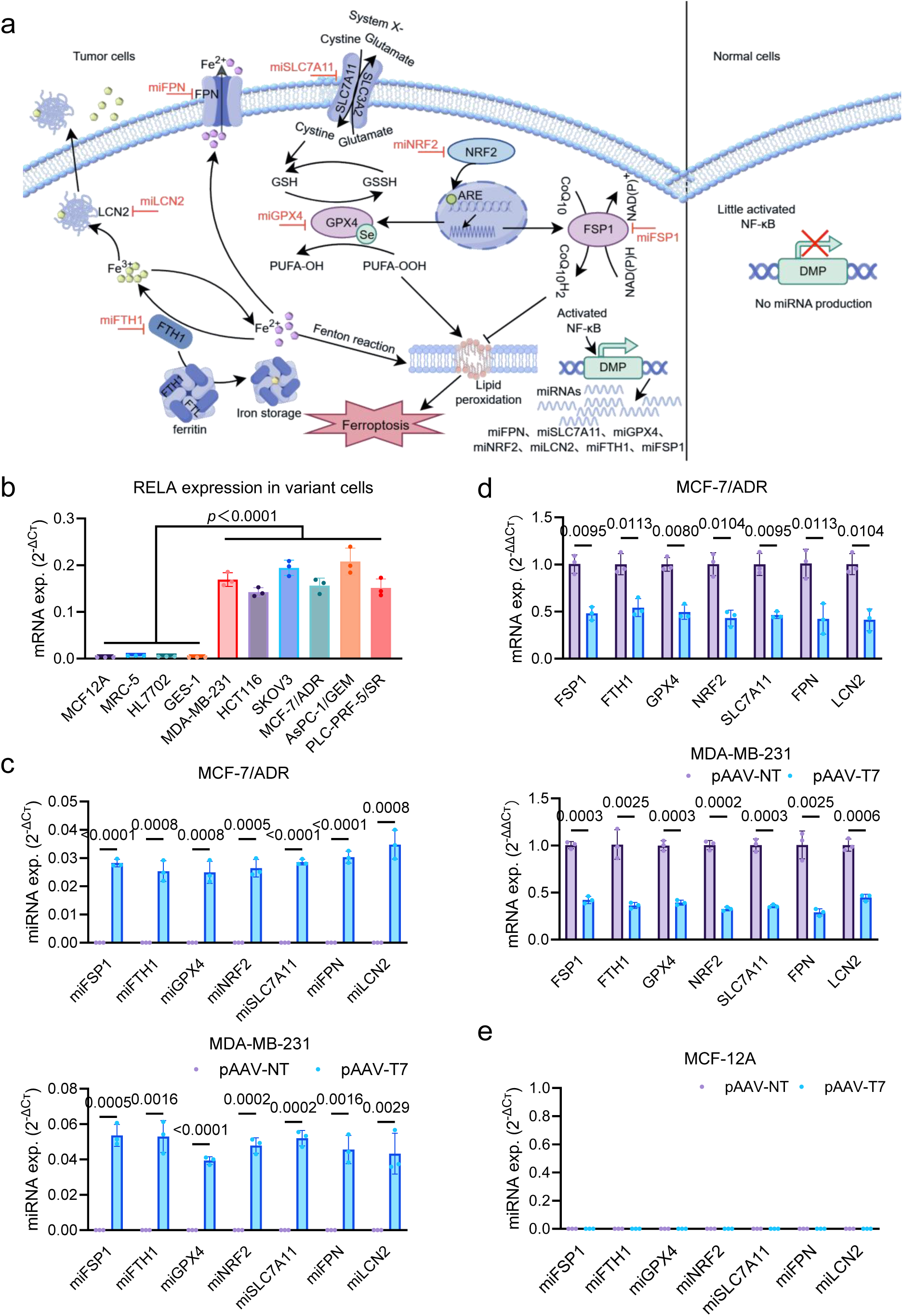
Construction and characterization of the NF-κB-responsive pAAV-T7 system. **a,** Schematic illustration of the pAAV-T7 system, in which the NF-κB-responsive promoter drives the expression of seven miRNAs targeting key regulators of ferroptosis, including miR-GPX4, miR-SLC7A11, miR-FSP1, miR-FPN, miR-NRF2, miR-LCN2, and miR-FTH1. **b,** RELA mRNA expression (exp.) levels in variant normal and cancer cells. **c,** Expression levels of miR-GPX4, miR-SLC7A11, miR-FSP1, miR-FPN, miR-NRF2, miR-LCN2, and miR-FTH1 in the MDA-MB-231 and MCF-7/ADR cells, respectively. **d,** Relative mRNA expression levels of the corresponding target genes including GPX4, SLC7A11, FSP1, FPN, NRF2, LCN2, and FTH1 in the MCF-7/ADR and MDA-MB-231 cells, respectively. **e,** Expression levels of miR-GPX4, miR-SLC7A11, miR-FSP1, miR-FPN, miR-NRF2, miR-LCN2, and miR-FTH1 in and in the MCF-12A cells. Data are shown as mean ± SD (n=3 wells). Statistical significance was analyzed by one-way ANOVA with Tukey’s test in (b), and by the unpaired two-tailed Student’s t-test in (c), (d) and (e). P values are given and that less than 0.05 means statistical significance.

### pAAV-T7 enhances chemotherapeutic sensitivity and reverses acquired chemoresistance in vitro

Having confirmed the tumor-selective expression of the miRNAs and the downregulation of their target genes involved in ferroptosis defense, we next investigated whether pAAV-T7 could enhance the sensitivity of tumor cells to chemotherapeutic agents and restore drug sensitivity in chemoresistant cells. Chemotherapy-resistant cancer cell lines (MCF-7/ADR and AsPC-1/GEM) and non-resistant tumor cell lines (MDA-MB-231, SKOV3, and HCT116) were transfected with either the empty pAAV-NT vector or pAAV-T7, followed by treatment with the corresponding chemotherapeutic agents (DOX, GEM, CDDP, or 5-FU) for 72 h. Cell viability was subsequently assessed using the CCK-8 assay.

As expected, treatment with the corresponding chemotherapeutic agents alone resulted in a dose-dependent decrease in cell viability. Notably, pAAV-T7 markedly reduced the IC_50_ values of the corresponding chemotherapeutic agents in both non-resistant and chemotherapy-resistant tumor cells. In chemotherapy-resistant cells, the IC_50_ values of DOX in MCF-7/ADR cells (13.14 μM vs. 0.5714 μM) and GEM in AsPC-1/GEM cells (76.86 μM vs. 4.443 μM) were significantly decreased following pAAV-T7 transfection (Fig. 2a, Fig. S2). Similarly, pAAV-T7 substantially increased the sensitivity of non-resistant MDA-MB-231 cells to DOX (3.547 μM vs. 0.1693 μM), SKOV3 cells to CDDP (8.264 μM vs. 2.496 μM), and HCT116 cells to 5-FU (8.080 μM vs. 1.382 μM) (Fig. 2a, Fig. S2). These findings indicate that pAAV-T7 enhances the response of tumor cells to chemotherapeutic agents and partially restores chemosensitivity in chemotherapy-resistant cells.

**Fig.2:**
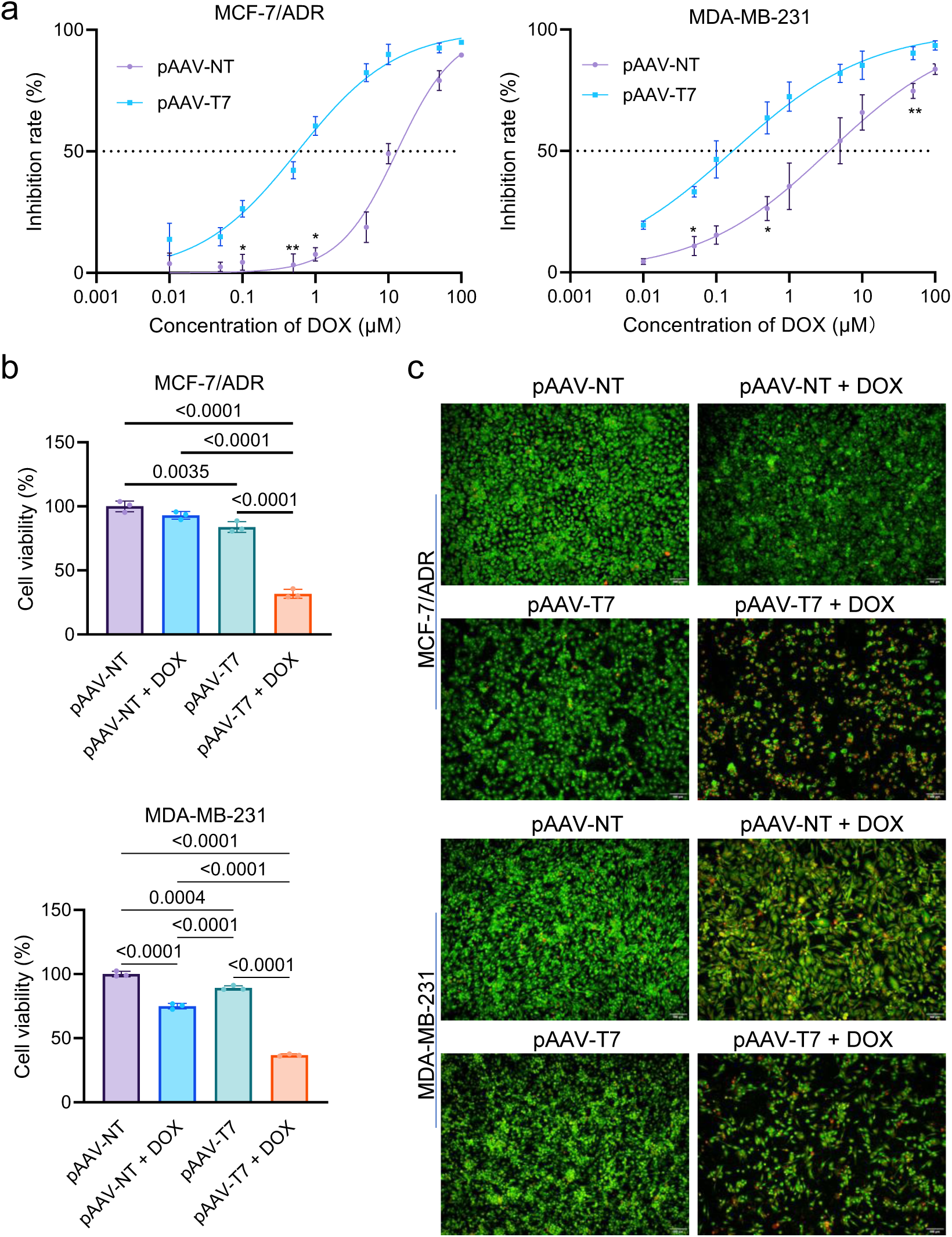
pAAV-T7 Enhances Chemotherapeutic Sensitivity and Reverses DOX Resistance. **a,** IC₅₀ values of DOX in MCF-7/ADR and MDA-MB-231 cells transfected with pAAV-NT or pAAV-T7. **b,** Cell viability of MCF-7/ADR and MDA-MB-231 cells following transfection with pAAV-NT or pAAV-T7 and subsequent treatment with DOX for 72 h, as determined by the CCK-8 assay. Data are shown as mean ± SD (n=3 wells). **c,** Representative fluorescence images of AO/EB staining of MCF-7/ADR and MDA-MB-231 cells transfected with pAAV-NT or pAAV-T7, with or without the corresponding anticancer agents for 72 h. MCF-7/ADR cells were treated with 1 μM DOX. MDA-MB-231 cells were treated with 0.5 μM DOX. Statistical significance was analyzed by one-way ANOVA with Tukey’s test in (c). IC_50_ values obtained from independent experiments were calculated by nonlinear regression using a four-parameter logistic model. IC_50_ values between the pAAV-NT and pAAV-T7 groups were compared using a two-tailed Student’s t-test. P values are given and that less than 0.05 means statistical significance.

Consistent with the IC_50_ analyses, CCK-8 assays further demonstrated that pAAV-T7 markedly enhanced the cytotoxic effects of the corresponding chemotherapeutic agents. Following drug exposure, pAAV-T7-transfected tumor cells exhibited a significantly greater reduction in cell viability than control cells, with this effect observed in both non-resistant and chemotherapy-resistant tumor cells (Fig. 2b, Fig. S3a). To further corroborate these findings, AO/EB staining was performed following combination treatment. pAAV-T7-transfected tumor cells displayed a markedly increased proportion of orange/red fluorescence-positive cells upon chemotherapeutic exposure, indicating enhanced cell death (Fig. 2c, Fig. S3b). Collectively, these findings demonstrate that pAAV-T7 enhances tumor-cell sensitivity to multiple chemotherapeutic agents and partially overcomes acquired chemoresistance in vitro.

### pAAV-T7 enhances chemotherapeutic sensitivity and reverses chemoresistance through ferroptosis-associated mechanisms

To elucidate the mechanism underlying pAAV-T7-mediated enhancement of chemotherapeutic sensitivity and reversal of chemoresistance, we first examined intracellular Fe²⁺ levels following combination treatment with pAAV-T7 and the corresponding chemotherapeutic agents. Combined treatment markedly increased intracellular Fe²⁺ accumulation, as demonstrated by fluorescence imaging and flow cytometry (Fig. 3a–c and Fig. S4a–c). We next assessed lipid peroxidation using C11-BODIPY 581/591. Combination treatment with pAAV-T7 and the corresponding chemotherapeutic agents induced a pronounced red-to-green fluorescence shift and significantly increased the 530/572 nm fluorescence ratio, indicating enhanced lipid peroxidation (Fig. 4a, b and Fig. S5a, b). Consistently, DCFH-DA staining revealed significantly elevated intracellular total ROS levels following combination treatment (Fig. 5a–c and Fig. S6a–c).

**Fig.3:**
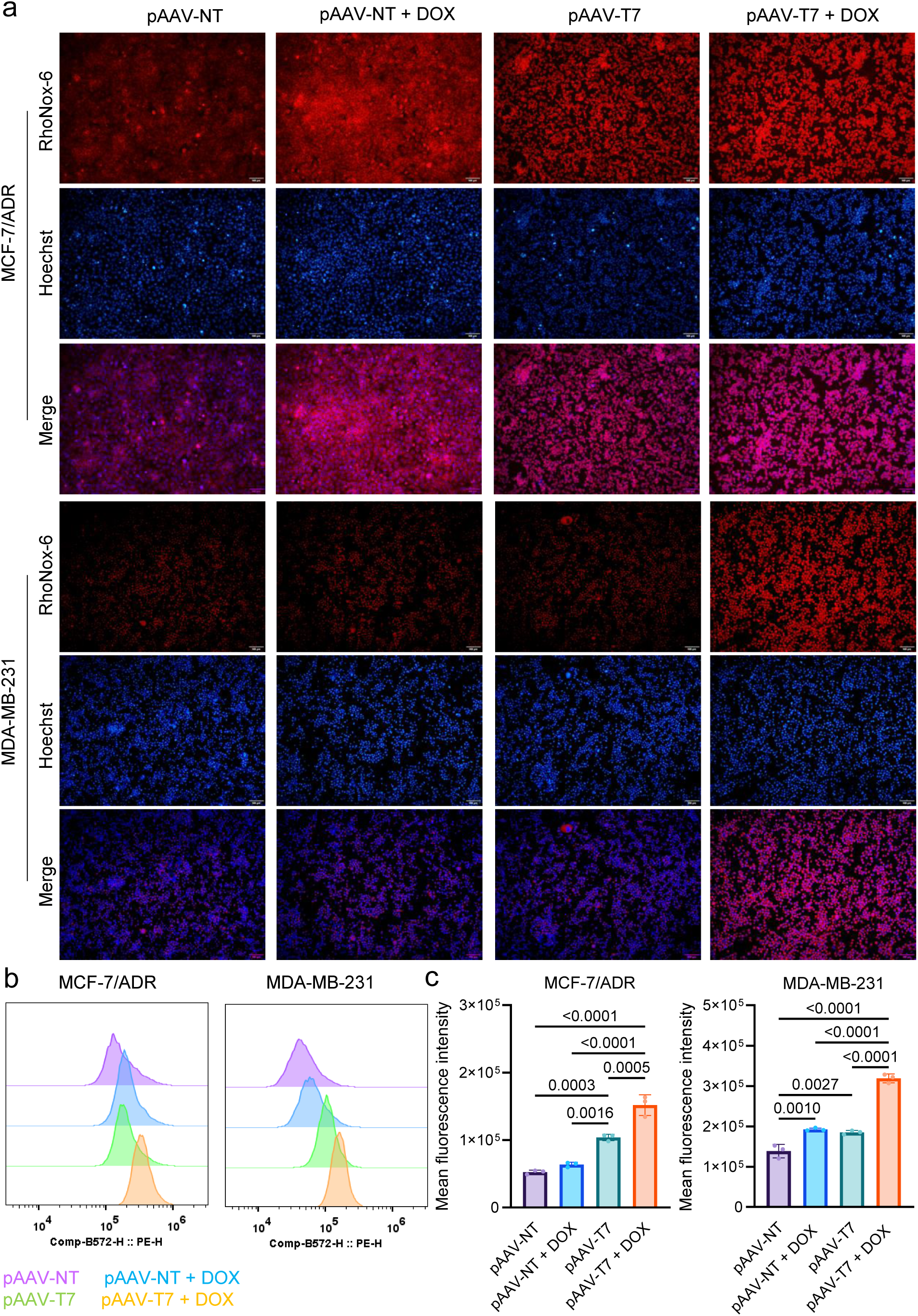
Combination of pAAV-T7 with DOX increases intracellular ferrous iron levels in tumor cells. MDA-MB-231 and MCF-7/ADR cells were transfected with pAAV-NT or pAAV-T7 and subsequently treated with DOX for 48 h. **a,** Representative fluorescence images of intracellular ferrous iron (Fe²⁺) staining. **b,** Representative flow-cytometry histograms showing intracellular Fe²⁺ levels. **c,** Quantification of the mean fluorescence intensity (MFI) of intracellular Fe²⁺ measured by flow cytometry. Data are shown as mean ± SD (n=3 independent experiments). Statistical significance was analyzed by one-way ANOVA with Tukey’s test in (c). P values are given and that less than 0.05 means statistical significance.

**Fig.4:**
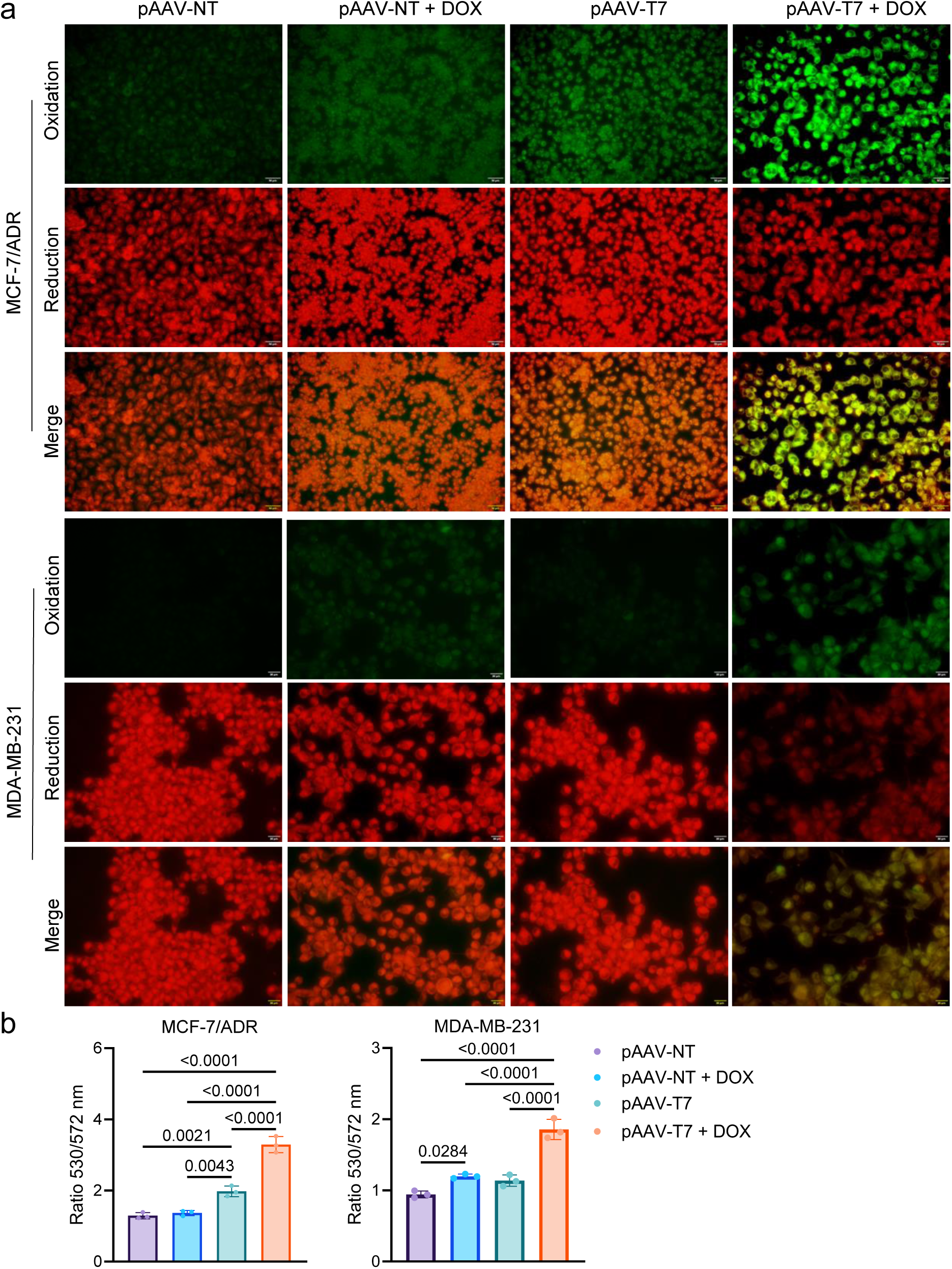
Combination of pAAV-T7 with DOX promotes lipid peroxidation in tumor cells. MDA-MB-231 and MCF-7/ADR cells were transfected with pAAV-NT or pAAV-T7 and subsequently treated with DOX for 48 h. **a,** Representative fluorescence images of BODIPY 581/591 C11 staining showing lipid peroxidation in MDA-MB-231 and MCF-7/ADR cells. **b,** Quantification of lipid peroxidation based on the fluorescence intensity of BODIPY 581/591 C11. Data are shown as mean ± SD (n=3 independent experiments). Statistical significance was analyzed by one-way ANOVA with Tukey’s test in (b). P values are given and that less than 0.05 means statistical significance.

**Fig.5:**
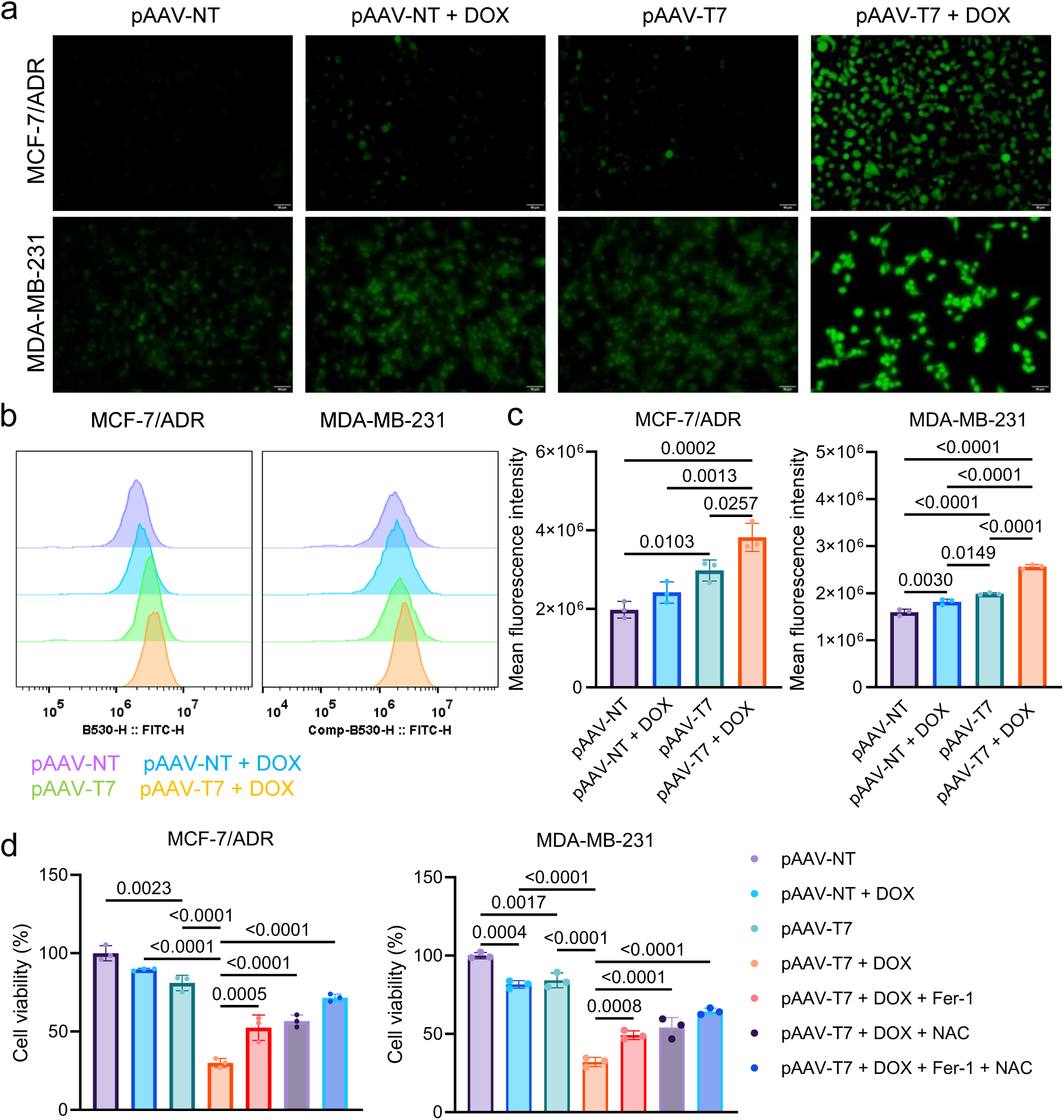
pAAV-T7 combined with anticancer agents increases intracellular ROS levels and induces ferroptosis in tumor cells. MDA-MB-231 and MCF-7/ADR cells were transfected with pAAV-NT or pAAV-T7 and subsequently treated with DOX. For ROS detection, cells were treated for 48 h. For the rescue experiments, cells were treated with DOX in combination with ferrostatin-1 (Fer-1) or/and N-acetylcysteine (NAC) for 72 h, followed by assessment of cell viability using the CCK-8 assay. **a,** Representative fluorescence images of DCFH-DA staining showing intracellular total ROS levels. **b,** Representative flow-cytometry histograms of DCFH-DA fluorescence. **c,** Quantification of the mean fluorescence intensity (MFI) of DCFH-DA fluorescence measured by flow cytometry. **d,** CCK-8-based rescue experiments showing the effects of Fer-1 and NAC on the cytotoxicity induced by the combination of pAAV-T7 and DOX. Data are shown as mean ± SD (n=3 independent experiments). Statistical significance was analyzed by one-way ANOVA with Tukey’s test in (c) and (d). P values are given and that less than 0.05 means statistical significance.

To further determine whether ferroptosis-associated processes and oxidative stress contributed to the enhanced cytotoxic effects of chemotherapy following pAAV-T7 treatment, we performed pharmacological rescue experiments using 1 μM ferrostatin-1 (Fer-1), a ferroptosis inhibitor, and 1 mM N-acetylcysteine (NAC), a ROS scavenger. In both tumor cell models, combined treatment with pAAV-T7 and DOX resulted in a pronounced reduction in cell viability. Addition of either Fer-1 or NAC significantly attenuated the cytotoxicity induced by the pAAV-T7/DOX combination. Notably, combined treatment with Fer-1 and NAC resulted in a further recovery of cell viability compared with either treatment alone (Fig. 5d and Fig. S7). Together, these findings indicate that ferroptosis-associated lipid peroxidation and oxidative stress contribute to the enhanced cytotoxic effects of chemotherapy induced by pAAV-T7.

### rAAV-T7 reverses chemoresistance in tumor spheroids and enhances chemotherapeutic sensitivity in patient-derived organoids

To further validate the chemotherapeutic sensitizing activity of rAAV-T7 in three-dimensional tumor models, we first established MCF-7/ADR tumor spheroids and evaluated their response to rAAV-T7 in combination with DOX. Calcein-AM/PI/Hoechst staining revealed a marked increase in PI-positive dead cells and a concomitant reduction in viable cells following combined treatment with rAAV-T7 and DOX (Fig. 6a). Consistently, 3D viability analysis demonstrated that rAAV-T7 significantly potentiated DOX-induced cytotoxicity, resulting in a pronounced reduction in tumor spheroid viability (Fig. 6b). Morphological analysis further showed that combination treatment substantially suppressed spheroid growth, as reflected by a significant reduction in mean spheroid diameter compared with the corresponding control groups (Fig. 6c). In addition, DCFH-DA staining revealed a marked increase in intracellular total ROS levels following combined treatment with rAAV-T7 and DOX (Fig. 6d), consistent with the enhanced oxidative stress observed in two-dimensional tumor cells.

**Fig.6:**
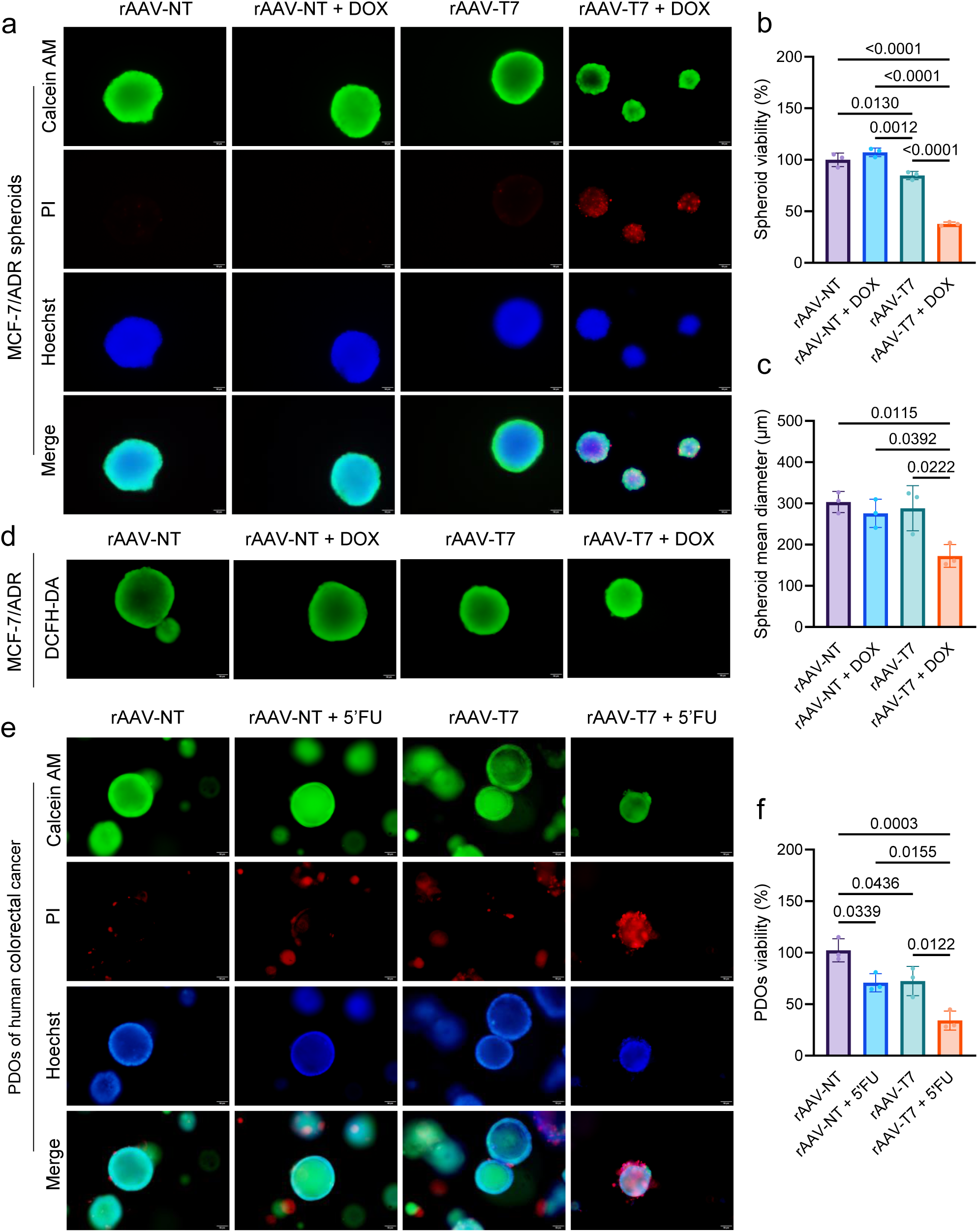
rAAV-T7 enhances the antitumor activity of anticancer agents in three-dimensional tumor models. MCF-7/ADR tumor spheroids were transduced with rAAV-NT or rAAV-T7 for 48 h and subsequently exposed to 1 μM doxorubicin (DOX) for 96 h. **a,** Representative fluorescence images of Calcein/PI/Hoechst live/dead staining of MCF-7/ADR tumor spheroids. **b,** Viability of MCF-7/ADR tumor spheroids determined using the Cell Counting Kit-3D assay. **c,** Quantification of the mean spheroid diameter. **d,** Representative fluorescence images of DCFH-DA staining showing intracellular ROS accumulation in MCF-7/ADR tumor spheroids. **e,** Representative fluorescence images of Calcein/PI/Hoechst live/dead staining of patient-derived organoids (PDOs) of human colorectal cancer. **f,** PDOs viability quantified using the Cell Counting Kit-3D assay. For PDOs of human colorectal cancer organoids, PDOs were transduced with rAAV-NT or rAAV-T7 for 12 h before being embedded in Matrigel. Following a 36h culture period, organoids were subsequently treated with 1 μM 5-fluorouracil (5-FU) for 96 h. Data are shown as mean ± SD (n=3 wells). Statistical significance was analyzed by one-way ANOVA with Tukey’s test in (b), (c) and (f). P values are given and that less than 0.05 means statistical significance.

We next investigated whether the chemotherapeutic sensitizing activity of rAAV-T7 could be extended to patient-derived tumor models. Colorectal cancer organoids (CRCOs) were treated with rAAV-T7 in combination with 5-FU and subsequently assessed by Calcein-AM/PI/Hoechst staining and 3D viability analysis. Combined treatment with rAAV-T7 and 5-FU resulted in a substantial increase in PI-positive dead cells and a reduction in viable organoid structures (Fig. 6e). Consistently, rAAV-T7 significantly enhanced the inhibitory effect of 5-FU on organoid viability (Fig. 6f). Notably, rAAV-T7 alone also made a similar significant antitumor effect in the PDOs. Collectively, these findings demonstrate that rAAV-T7 enhances the efficacy of chemotherapeutic agents in three-dimensional tumor models, reverses DOX resistance in MCF-7/ADR tumor spheroids, and increases 5-FU sensitivity in CRCOs.

### rAAV-T7 Enhances Chemotherapeutic Sensitivity and Reverses DOX Resistance In Vivo

To evaluate the in vivo antitumor efficacy and tolerability of rAAV-T7, subcutaneous xenograft models of MCF-7/ADR tumor were established in nude mice. When tumor volumes reached approximately 150 mm³, mice were randomly allocated to four groups: rAAV-NT, rAAV-NT + DOX, rAAV-T7, and rAAV-T7 + DOX. Virus was administered intravenously at 1.5 × 10_11_ vg/mouse, while DOX was administered intraperitoneally at 2 mg/kg (Fig.7a). Tumor growth and body weight were monitored throughout the treatment period. Tumor growth curve, tumor imaging and tumor volume and weight at the endpoint showed that rAAV-NT+DOX did not significantly inhibit tumor growth in comparison with rAAV-NT, consistent with the DOX-resistant phenotype of MCF-7/ADR cells (Fig.7b–d). However, in comparison with rAAV-NT, rAAV-T7 alone significantly inhibit tumor growth (Fig.7b–d). Importantly, the combined treatment with rAAV-T7 and DOX obtained the most significant antitumor effect, superior to rAAV-T7 alone, suggesting rAAV-T7 in the combined treatment resensitizes tumors to DOX and made a synergistic antitumor effect.

To further evaluate the in vivo antitumor efficacy and tolerability of rAAV-T7, subcutaneous xenograft models of MDA-MB-231 tumors were then established in NCG mice. Similarly, when tumor volumes reached approximately 150 mm³, mice were randomly allocated to the same four groups: rAAV-NT, rAAV-NT + DOX, rAAV-T7, and rAAV-T7 + DOX. Virus was administered intravenously at 1.5 × 10^11^ vg/mouse, while DOX was administered intraperitoneally at 1 mg/kg (Fig.7e). Tumor growth and body weight were also monitored throughout the treatment period. As evidenced by tumor growth curve (Fig. 7f), and tumor imaging (Fig. 7g) and tumor volume and weight (Fig. 7h) at the endpoint, a similar therapeutic response was observed in MDA-MB-231 tumor-bearing NCG mice. rAAV-NT+DOX produced a significant inhibition of tumor growth in comparison with rAAV-NT alone, consistent with the important role of chemotherapy in the treatment of triple-negative breast cancer (TNBC). Notably, rAAV-T7 alone also obtained the similar antitumor effect to rAAV-NT+DOX. Importantly, compared with the two monotherapies, the combination of rAAV-T7 and DOX resulted in the most prominent tumor growth suppression, superior to either monotherapy, also suggesting that rAAV-T7 make tumors more sensitive to DOX and made a synergistic antitumor effect together with DOX.

**Fig.7:**
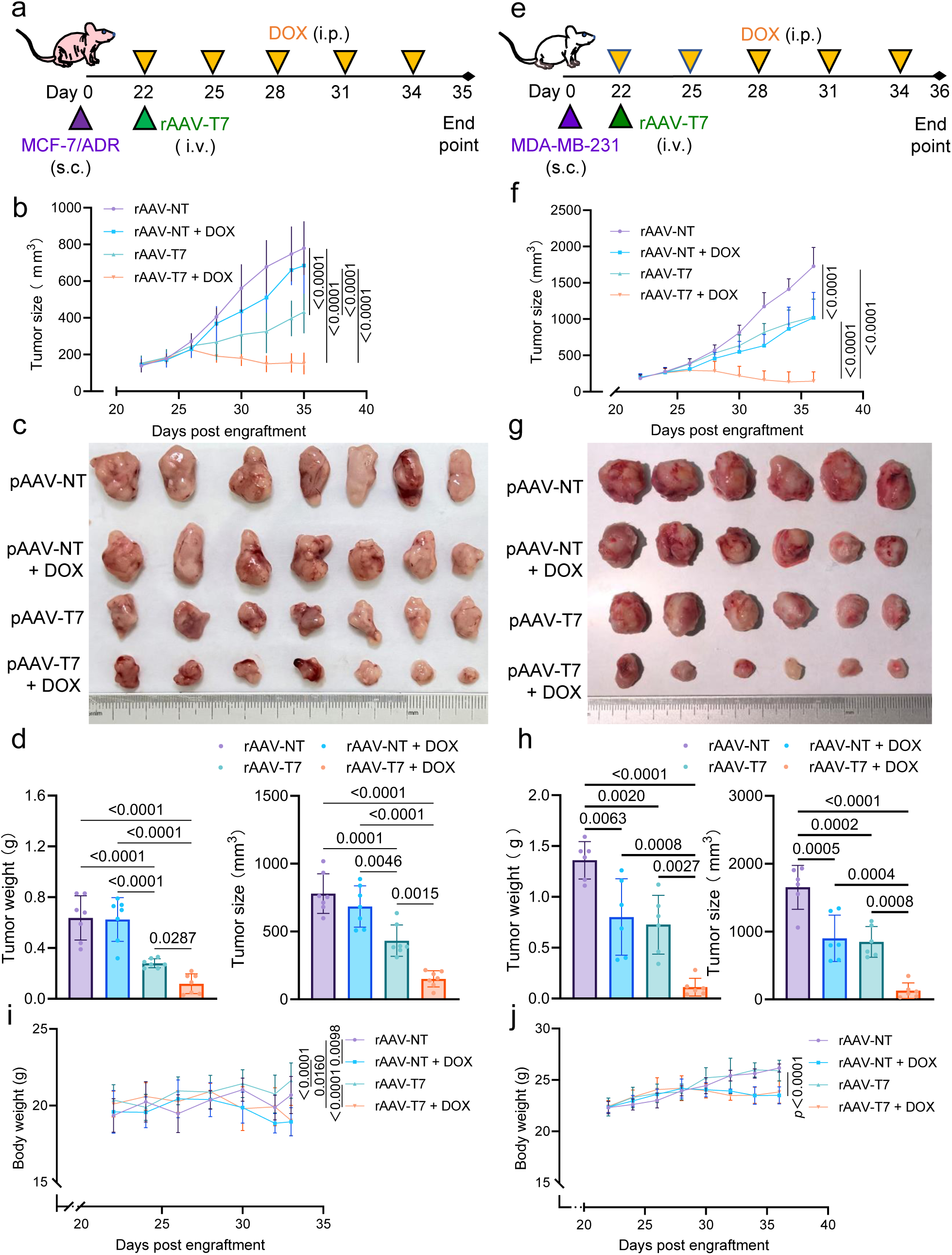
rAAV-T7 enhances the antitumor efficacy of DOX in vivo. **a,** Schematic illustration of the treatment schedule for mice bearing MCF-7/ADR xenograft tumors. **b,** Tumor growth curves of MCF-7/ADR xenografts during the treatment period (n=7). **c,** Photographs of excised MCF-7/ADR xenograft tumors at the endpoint of treatment (n=7). **d,** Tumor weights and tumor sizes of excised MCF-7/ADR xenografts at the endpoint of treatment (n=7). **e,** Schematic illustration of the treatment schedule for mice bearing MDA-MB-231 xenograft tumors. **f,** Tumor growth curves of MDA-MB-231 xenografts during the treatment period (n=6). **g,** Photographs of excised MDA-MB-231 xenograft tumors at the endpoint of treatment (n=6). **h,** Tumor weights and tumor sizes of excised MDA-MB-231 xenografts at the endpoint of treatment (n=6). **i,** Body weight changes of MCF-7/ADR tumor-bearing mice during treatment (n=7). **j,** Body weight changes of MDA-MB-231 tumor-bearing mice during treatment (n=6). Data are shown as mean ± SD. Statistical significance was analyzed by one-way ANOVA with Tukey’s test in (b), (d), (f), (h), (i), and (j). P values are given and that less than 0.05 means statistical significance.

Body weight was monitored as an indicator of general health and treatment tolerability. Although the DOX-treated groups exhibited lower body weights than the corresponding non-DOX groups at the treatment endpoint, no significant body weight loss was observed relative to baseline in any treatment group. Moreover, body weight did not differ significantly between the rAAV-T7+DOX and rAAV-NT+DOX groups, indicating that rAAV-T7 did not exacerbate DOX-associated body weight changes (Fig. 7i, j).

We next evaluated the systemic safety of the treatment. In the MCF-7/ADR tumor-bearing nude mice, H&E staining of the heart, liver, spleen, lung, and kidney revealed no obvious pathological abnormalities (Fig. S8a). Serum ALT, AST, ALP, BUN, Cr, and UA levels remained within their respective normal reference ranges, with no significant differences among the treatment groups (Fig. S8b). Although serum CK-MB levels were elevated in DOX-treated mice, no significant difference was observed between the rAAV-NT+DOX and rAAV-T7+DOX groups (Fig. S8b), suggesting that rAAV-T7 did not exacerbate DOX-associated cardiac toxicity. In the MDA-MB-231 tumor-bearing NCG mice, similar results of H&E staining (Fig. S9a), serum AST, ALP, BUN, Cr, and UA levels (Fig. S9b) were observed. Although serum ALT levels were significantly elevated in both DOX-treated groups relative to the corresponding non-DOX groups, they remained within the normal reference range, suggesting a mild DOX-associated hepatic response rather than overt hepatotoxicity. Importantly, ALT levels did not differ significantly between the rAAV-NT+DOX and rAAV-T7+DOX groups (Fig. S9b), indicating that rAAV-T7 did not exacerbate DOX-associated hepatic effects. Likewise, although serum CK-MB levels were elevated in DOX-treated mice, no significant difference was observed between the rAAV-NT+DOX and rAAV-T7+DOX groups (Fig. S9b), further suggesting that rAAV-T7 did not aggravate DOX-associated cardiac effects. Collectively, these findings suggest that rAAV-T7 did not cause overt pathological damage to major organs or exacerbate the systemic effects associated with DOX treatment.

To further investigate the tumor selectivity of rAAV-T7 activity in vivo, we examined the biodistribution of rAAV vectors and the expression of the encoded miRNAs and their corresponding target genes in major organs and tumor tissues. In the MCF-7/ADR tumor-bearing nude mice, qPCR analysis revealed that rAAV vector genomes were detectable in multiple tissues, including the heart, liver, spleen, lung, and kidney, indicating that the vectors were distributed systemically following intravenous administration (Fig. S10a). However, RELA mRNA expression was substantially higher in tumor tissues than in the examined normal organs (Fig. S10b), suggesting enhanced NF-κB activity in the tumor tissues. Accordingly, despite the broad rAAV tissue distribution, all seven rAAV-T7-encoded miRNAs, including miGPX4, miSLC7A11, miFSP1, miFPN, miNRF2, miLCN2, and miFTH1, were detected only in tumor tissues, but not in the examined normal organs (Fig. S10c). Consistently, the corresponding target genes, GPX4, SLC7A11, FSP1, FPN, NRF2, LCN2, and FTH1, were significantly downregulated in tumor tissues, whereas their expression levels remained largely unchanged in the heart, liver, spleen, lung, and kidney (Fig. S11). In MDA-MB-231 tumor-bearing NCG mice, similar results of rAAV vector genome distribution (Fig. S12a), RELA mRNA expression (Fig. S12b), miRNA expression (Fig. S12c), and miRNA target gene expression (Fig. S13) were observed. Collectively, these findings demonstrate that although rAAV vectors are distributed across multiple tissues, rAAV-T7-mediated miRNA expression and target gene suppression are specifically restricted to tumor tissues, potentially owing to the elevated NF-κB activity within the tumor microenvironment.

Finally, to determine whether rAAV-T7/DOX treatment induced ferroptosis-associated alterations in vivo, we measured Fe²⁺, MDA, and GSH levels in MDA-MB-231 xenograft tissues. Compared with either monotherapy, combined treatment with rAAV-T7 and DOX significantly increased intratumoral Fe²⁺ and MDA levels while reducing GSH levels (Fig. S14). These changes were consistent with the ferroptosis-associated alterations observed in vitro, further supporting the involvement of ferroptosis in the enhanced antitumor activity of the rAAV-T7/DOX combination. Collectively, these findings demonstrate that rAAV-T7 specifically induces the expression of seven miRNAs in tumor tissues, resulting in downregulation of their corresponding target genes and promotion of ferroptosis-associated metabolic alterations. This tumor-selective activity potentiates the antitumor efficacy of DOX while producing no additional toxicity in major organs.

### rAAV-T7 Enhances Sorafenib Sensitivity in Sorafenib-Resistant Tumor Cells

Although chemotherapy represents a major component of systemic cancer treatment, targeted therapies are also widely used in clinical practice, and acquired resistance to targeted agents remains a major challenge that limits their therapeutic efficacy. We therefore sought to determine whether the ferroptosis-sensitizing activity of rAAV-T7 could be extended beyond chemotherapy to overcome resistance to targeted therapy, using sorafenib-resistant PLC-PRF-5/SR cells as a model. We first assessed the dose–response of PLC-PRF-5/SR cells to sorafenib following pAAV-T7 transfection. pAAV-T7 markedly enhanced sorafenib sensitivity, reducing the sorafenib IC_50_ from 15.28 μM to 2.341 μM (Fig. S15a). Consistently, CCK-8 assays showed that combined treatment with pAAV-T7 and sorafenib significantly reduced cell viability compared with either treatment alone (Fig. S15b). AO/EB staining further confirmed the enhanced cytotoxic effect of the combination treatment, with a markedly increased proportion of orange/red fluorescence-positive cells (Fig. S15c).

We next examined ferroptosis-associated responses following combination treatment. Combined treatment with pAAV-T7 and sorafenib markedly increased intracellular Fe²⁺ accumulation, as determined by fluorescence imaging and flow cytometry (Fig. S16a-c). C11-BODIPY 581/591 staining further revealed enhanced lipid peroxidation, reflected by an increased 530/572 nm fluorescence ratio (Fig. S17a, b). In addition, DCFH-DA analysis demonstrated a significant increase in intracellular total ROS levels following combination treatment (Fig. S17c-e). Collectively, these findings demonstrate that pAAV-T7 sensitizes sorafenib-resistant PLC-PRF-5/SR cells to sorafenib and is associated with increased Fe²⁺ accumulation, lipid peroxidation, and oxidative stress.

## Discussion

Breast cancer is the most commonly diagnosed malignancy in women worldwide. Triple-negative breast cancer (TNBC), in particular, is characterized by aggressive behavior, a high risk of recurrence, and poor prognosis^22^. Although chemotherapy remains a cornerstone of systemic treatment for TNBC^23^, resistance to anthracyclines and taxanes can emerge during treatment and compromise therapeutic efficacy^24^. Pancreatic cancer is among the most lethal malignancies, with aggressive progression, limited treatment options, and poor clinical outcomes. Its therapeutic management is further complicated by substantial resistance to chemotherapy, including GEM-based regimens^25^. Liver cancer also represents a major global health burden. Sorafenib has been an established systemic treatment for advanced hepatocellular carcinoma, but its clinical benefit is limited in many patients^26^, in part because of intrinsic or acquired resistance^27^. These challenges underscore the need for strategies that can improve tumor responsiveness to both conventional chemotherapy and targeted therapy.

Cancer cells frequently alter iron metabolism and redox homeostasis to support their high growth. Tumors therefore sustain fragile iron and redox homeostasis at a high level^28^. These changes create cancer cells a vulnerability to ferroptosis, an iron-dependent form of regulated cell death driven by lipid peroxidation^29^. Cancer cells in a high mesenchymal therapy-resistant cell state are dependent on the lipid hydroperoxidase GPX4 for survival^30^. Ferroptosis inducers or GPX4 targeting can induces ferroptosis in these resistant cells^31,32^, and effectively suppresses tumor relapse in mice^33^. Ferroptosis has thus emerged as a potential strategy for improving treatment responses in some drug-resistant tumors^8,12^. However, the current ferroptosis inducers, especially those of small molecules, generally lack intrinsic tumor selectivity, limiting their clinical application. Achieving selective ferroptosis in tumor cells while minimizing damage to normal tissues remains a major challenge. To overcome the bottleneck, we developed a rAAV-T7-based smart ferroptosis, which can sense the elevated NF-κB activity in tumor cells and then allow ferroptosis to just occur in tumor cells.

The cancer cell specificity of the smart ferroptosis is dependent on the activity of NF-κB. NF-κB signaling is frequently aberrantly or constitutively activated in human cancers and contributes to tumor-cell survival, proliferation, metabolism, and therapy resistance^34,35^. Consistent with this biological feature, RELA expression was higher in tumor cells and tissues than in normal cells and tissues (Fig.1b, Fig.S10, and Fig.S12). In animals, although rAAV-T7 vector genomes were detected in multiple organs following systemic administration (Fig.S10 and Fig.S12), the encoded miRNAs were only expressed in tumor tissues (Fig.S10 and Fig.S12), and suppression of their target genes were only observed in tumor tissues (Fig.S11 and Fig.S13). These findings reveal the high NF-κB-dependent tumor selectivity of this smart ferroptosis. In consistence, this smart ferroptosis showed biosafety in animals, indicated by monitoring of body weight, histopathology, and serum biochemistry. This design may therefore address an important limitation of systemic ferroptosis-inducing strategies, for which insufficient tumor selectivity and potential damage to normal tissues remain major concerns for therapeutic translation^9,13^.

Increasing evidence indicates cells resist ferroptosis at multiple pathways, such as GPX4–GSH, FSP1– CoQ₁₀–NAD(P)H, GCH1–BH₄, and DHODH–CoQH₂^36,37^. Disrupting individuals of these pathways with the current ferroptosis inducers can make cancer cells sensitive to ferroptosis and enhance their sensitivity to anticancer therapies^38–42^. However, cancer cells can offset the sensitivity by strengthening other redundant anti-ferroptosis pathways^43^. Therefore, rather than targeting a single anti-ferroptosis target, rAAV-T7 simultaneously targets seven key anti-ferroptosis targets including GPX4, SLC7A11, FSP1, NRF2, FPN, LCN2, and FTH1, which collectively contribute to antioxidant defense, lipid peroxide detoxification, and iron homeostasis sustainment^15,31,40,44–48^. This multiple-pathways co-targeting strategy make cancer cells unable to easily evade from the smart ferroptosis. Therefore, pAAV-T7 enhanced the effects of DOX, GEM, CDDP, and 5-FU across different tumor models and restored DOX sensitivity in MCF-7/ADR and GEM sensitivity in AsPC-1/GEM cells (Fig.2a, Fig.S2). Consistently, neither Fer-1 nor NAC alone, nor their combination, fully rescued cell viability following combined pAAV-T7 and chemotherapy treatment (Fig.5d). This incomplete rescue supports the rationale for simultaneously targeting multiple ferroptosis defense pathways, which may limit compensatory responses and allow tumor cells more sensitive to these treatments. Together, these findings suggest that the smart ferroptosis sensitizes resistant tumor cells to multiple chemotherapeutic agents by co-weakening multiple ferroptosis-resistant pathways.

Molecular-targeted therapies are widely used in cancer treatment, but therapeutic resistance remains a major clinical challenge^2^. Ferroptosis has emerged as a potential strategy for addressing this problem by exploiting vulnerabilities in tumor-cell redox regulation and ferroptosis defense^9^. Recent studies have identified GPX4-centered regulatory mechanisms as important components of ferroptosis resistance in HCC and have shown that disrupting these defenses can enhance the activity of molecular-targeted therapies^49–51^. Building on these findings, the present study simultaneously targets multiple ferroptosis-defense and iron-regulatory pathways. Although the contribution of individual pAAV-T7 targets may vary across tumor types and resistance states, their coordinated suppression may enhance tumor-cell sensitivity to anticancer treatments. Consistent with this possibility, pAAV-T7 enhanced the response to sorafenib in sorafenib-resistant PLC-PRF-5/SR cells and was accompanied by increased Fe²⁺ accumulation, lipid peroxidation, and oxidative stress. These findings suggest that coordinated disruption of ferroptosis and redox defenses may increase the vulnerability of resistant tumor cells to targeted therapy.

Notably, rAAV-T7 alone significantly inhibited tumor growth in our breast cancer models in vivo. This effect may be related to the disruption of multiple ferroptosis-defense pathways, which could render tumor cells more vulnerable to endogenous iron-dependent lipid peroxidation. Clinical studies have reported higher total iron concentrations in breast tumors than in adjacent non-tumor tissues^52^, while recent work has demonstrated that targeted GPX4 degradation can suppress breast cancer growth by promoting ferroptosis^53^. These findings provide a rationale for exploring rAAV-T7 monotherapy in tumors with elevated iron levels, where disruption of ferroptosis-defense pathways may increase susceptibility to endogenous iron-dependent lipid peroxidation. Unlike conventional ferroptosis inducers, the NF-κB-responsive design of rAAV-T7 may favor miRNA expression in tumor cells with elevated NF-κB activity, potentially improving tumor selectivity and reducing effects on normal tissues. This approach may therefore offer a potentially safer strategy for targeting iron-rich tumors, although its safety advantage over conventional ferroptosis inducers requires further validation.

Despite the encouraging therapeutic efficacy and tumor-selective activity observed in our study, several limitations should be considered. First, the current in vivo studies were primarily performed in immunodeficient models and therefore do not fully recapitulate the interactions among ferroptosis, NF-κB signaling, AAV immunity, and the tumor immune microenvironment. Second, although no detectable additional toxicity was observed in major organs with the treatment regimen used, vector immunogenicity, repeated administration, long-term vector persistence, and long-term safety remain to be evaluated before clinical translation. Third, the current clinical therapies may induce NF-κB activation in normal tissues and increase rAAV-T7 expression outside tumors and reduce its tumor selectivity. Nevertheless, in our animal studies, DOX treatment did not significantly alter RELA expression in the examined normal tissues. In consistence, we detected no significant miRNA expression or suppression of the target genes in these tissues. These findings suggest that, under the dosing regimens used in this study, NF-κB-responsive rAAV-T7 expression remained limited in the examined normal tissues. Although CK-MB levels increased following DOX treatment, the addition of rAAV-T7 did not further increase CK-MB levels compared with DOX treatment alone. This finding provides preliminary evidence that rAAV-T7 did not exacerbate this indicator of DOX-associated cardiac injury under the conditions tested. Future studies should therefore systematically evaluate the relationship between drug dose, treatment schedule, tissue-specific NF-κB activation, and rAAV-T7 transgene expression, particularly under repeated-treatment conditions.

## Conclusion

In conclusion, we developed an NF-κB-responsive AAV-T7 platform for tumor-selective sensitization to anticancer therapies by targeting ferroptosis and redox defenses. AAV-T7 specifically induced the expression of seven miRNAs in tumor cells, thereby weakening ferroptosis-defense mechanisms and increasing Fe²⁺ accumulation, lipid peroxidation, and oxidative stress. This multi-target strategy enhanced the efficacy of chemotherapeutic agents and the targeted agent sorafenib across multiple tumor models and increased the sensitivity of resistant tumor cells to treatment. Importantly, AAV-T7-mediated miRNA expression and target-gene suppression were only observed in tumor tissues and were not accompanied by detectable additional toxicity in major organs. Collectively, these findings support AAV-T7 as a smart ferroptosis inducer with tumor selectivity and capability to enhance the sensitivity of resistant tumors to anticancer therapies, which provides a new tool for overcoming resistance of tumors to those first-line drugs.

## Materials and methods

### Cell culture

HEK-293T (human embryonic kidney), HCT116 (human colorectal cancer), MDA-MB-231 (human triple-negative breast cancer), SKOV3 (human ovarian cancer), MCF-12A (human mammary epithelial), and GES-1 (human gastric epithelial) cells were cultured in Dulbecco’s modified Eagle’s medium (DMEM) supplemented with 10% fetal bovine serum (FBS), 100 U/mL penicillin, and 100 μg/mL streptomycin. MRC-5 (human fetal lung fibroblast) cells were maintained in minimum essential medium (MEM) supplemented with 10% FBS, 1% non-essential amino acids (NEAA), 100 U/mL penicillin, and 100 μg/mL streptomycin. HL7702 (human normal hepatocyte) cells were cultured in Roswell Park Memorial Institute 1640 medium (RPMI-1640) supplemented with 10% FBS, 100 U/mL penicillin, and 100 μg/mL streptomycin. MCF-7/ADR (human doxorubicin-resistant breast cancer) cells were maintained in RPMI-1640 supplemented with 10% FBS, 100 U/mL penicillin, and 100 μg/mL streptomycin, with 500 ng/mL doxorubicin (DOX) added to maintain the drug-resistant phenotype. AsPC-1/GEM (human gemcitabine-resistant pancreatic cancer) cells were cultured in DMEM supplemented with 10% FBS, 100 U/mL penicillin, and 100 μg/mL streptomycin, with 18 μg/mL gemcitabine (GEM) added to maintain gemcitabine resistance. PLC-PRF-5/SR (human sorafenib-resistant hepatocellular carcinoma) cells were maintained in MEM supplemented with 10% FBS, 100 U/mL penicillin, and 100 μg/mL streptomycin, with 2 μM sorafenib added to maintain sorafenib resistance. MCF-7/ADR, AsPC-1/GEM, and PLC-PRF-5/SR cells were cultured in drug-free medium for 2 weeks before the corresponding experiments. All cells were maintained at 37°C in a humidified atmosphere containing 5% CO₂.

### Cell transfection and Drug Treatment

Cells were transfected with plasmid DNA using different transfection reagents according to the cell type and experimental requirements. ExFect Transfection Reagent (Vazyme Biotech, T101) and Lipofectamine™ 3000 Transfection Reagent (Thermo Fisher Scientific, L3000015) were used for plasmid transfection. Transfection was performed according to the manufacturer’s instructions. Briefly, before transfection, the culture medium was replaced with serum-free and antibiotic-free medium. Plasmid DNA was mixed thoroughly with the corresponding transfection reagent according to the manufacturer’s instructions and subsequently added to the cells. After 12 h of transfection, the medium was replaced with complete culture medium. MCF-7/ADR and MDA-MB-231 cells were subsequently treated with 1 μM and 0.5 μM DOX, respectively; SKOV3 cells were treated with 2 μM cisplatin (CDDP); AsPC-1/GEM cells were treated with 10 μM GEM; HCT116 cells were treated with 1 μM 5-fluorouracil (5-FU); and PLC-PRF-5/SR cells were treated with 2 μM sorafenib. Cells without drug treatment were used as corresponding controls. After 48 h of treatment, intracellular Fe²⁺ levels, lipid peroxidation, and ROS levels were analyzed. After 72 h of treatment, cell death and cell viability were evaluated by AO/EB staining and the corresponding cell viability assay, respectively.

### Virus preparation

rAAV vectors were produced and purified as previously described^19^. Briefly, HEK-293T cells were co-transfected with the corresponding pAAV transfer plasmid, pHelper, and pAAV-RC plasmids using ExFect Transfection Reagent. After 72 h, rAAV vectors were harvested and purified by freeze–thaw lysis, chloroform extraction, and polyethylene glycol 8000 (PEG 8000) precipitation. Vector genome titers were determined by quantitative polymerase chain reaction (qPCR), and purified rAAV vectors were aliquoted and stored at −80°C until use.

### Experiments in Tumor Spheroids

MCF-7/ADR (human doxorubicin-resistant breast cancer) cells were cultured in Dulbecco’s modified Eagle’s medium (DMEM) supplemented with B-27™ Supplement (Gibco, 10889-038), N-2 Supplement (Gibco, 17502-048), 20 ng/mL epidermal growth factor (EGF; Thermo Fisher Scientific, PHG0314), and 20 ng/mL basic fibroblast growth factor (bFGF; R&D Systems, 3718-FB-010). Cells were seeded into 96-well plates at a density of 1,000 cells per well and cultured until tumor spheroids formed. Cells were seeded into 96-well plates at a density of 1,000 cells per well and cultured until tumor spheroids formed. Each well containing a tumor spheroid was then treated with 1 × 10⁸ vector genomes (vg) of rAAV-NT or rAAV-T7 and cultured for an additional 48 h. Subsequently, spheroids were treated with or without 1 μM doxorubicin (DOX) for 96 h. Spheroid diameter was measured from images using ImageJ software. Reactive oxygen species (ROS) levels were detected using the Beyo3D™ Reactive Oxygen Species Assay Kit (DCFH-DA) (Beyotime, S1105S) according to the manufacturer’s instructions. Spheroid viability was evaluated using the Beyo3D™ Calcein/PI/Hoechst Cell Viability Assay Kit (Beyotime, C1375S), and overall spheroid viability was quantified using the Cell Counting Kit-3D (Beyotime, C0049S) according to the manufacturer’s instructions.

### Experiments in PDO

The culture medium was removed, and the organoids were treated with 2 mL of TrypLE™ Express (Gibco, 12604013) at 37°C for 5 min. The organoids were gently pipetted to disperse the Matrigel and then placed on a shaker for 5 min. Subsequently, 2 mL of culture medium was added, and the suspension was gently pipetted to achieve a homogeneous cell suspension. The cells were centrifuged at 1,200 rpm for 5 min at 4°C, and the supernatant was discarded. The cell pellet was washed once with culture medium and resuspended in 1 mL of culture medium for cell counting. Cells were seeded into low-attachment 24-well plates at a density of 1 × 10⁵ cells per well in 500 μL of colorectal cancer organoid culture medium (bioGenous, K2103-CR) and cultured for 12 h. Each well was then treated with 1 × 10¹⁰ vector genomes (vg) of rAAV-NT or rAAV-T7 and mixed thoroughly, followed by an additional 48 h of culture. The cell culture suspension containing organoids was collected and centrifuged at 300 × g for 5 min. After removal of the supernatant, the organoids were washed once with organoid passaging buffer and resuspended in 100 μL of Matrigel (bioGenous, M315066). A 5-μL aliquot of the Matrigel-organoid mixture was seeded into a 96-well plate and incubated at 37°C for 15 min to allow Matrigel polymerization. Subsequently, 100 μL of culture medium was added to each well, and the organoids were cultured for 48 h, followed by treatment with or without 1 μM 5-FU for an additional 96 h. Organoid viability was evaluated using the Beyo3D™ Calcein/PI/Hoechst Cell Viability Assay Kit (Beyotime, C1375S), and overall organoid viability was quantified using the Cell Counting Kit-3D (Beyotime, C0049S) according to the manufacturer’s instructions.

### Quantitative PCR

Total RNA was extracted from cell lines and tissue samples using RNAiso Plus (Takara, 9108) according to the manufacturer’s instructions. Complementary DNA (cDNA) was synthesized using the PrimeScript™ RT reagent Kit with gDNA Eraser (Takara, RR047A) according to the manufacturer’s instructions. Genomic DNA (gDNA) was isolated from tissue samples using the FastPure Cell/Tissue DNA Isolation Mini Kit (Vazyme, DC102-01). Quantitative PCR (qPCR) was performed using an ABI StepOnePlus Real-Time PCR System (Applied Biosystems) with TB Green® Premix Ex Taq™ II (Tli RNaseH Plus) (Takara, RR820A). Each sample was analyzed in triplicate, and all experiments were independently repeated at least three times. For mRNA analysis, expression levels were calculated using the 2^-ΔCt^ or 2^-ΔΔCt^ method, where ΔCt = Ct_target_-Ct_GAPDH_ and ΔΔCt = ΔCt_treated_-ΔCt_control_. Viral DNA abundance was normalized to GAPDH and calculated using the 2^-ΔCt^ method. miRNA expression levels were normalized to U6 and calculated using the 2^-ΔCt^ method. Primer specificity was confirmed by melting curve analysis. All primer sequences are listed in Supplementary Table 1 and Supplementary Table 2.

### AO&EB Staining

Cell death was evaluated by acridine orange/ethidium bromide (AO/EB) staining using an AO/EB staining kit (Sangon Biotech, E607308). Cells were seeded into 96-well plates and cultured until they reached approximately 70–80% confluence. Cells were then transfected with the indicated plasmids and subsequently treated with the corresponding anticancer agents as described above. After 72 h of drug treatment, the cells were gently washed three times with phosphate-buffered saline (PBS). A staining solution was prepared by mixing 90 μL of Buffer with 5 μL each of AO and EB staining solutions. Subsequently, 100 μL of the freshly prepared staining solution was added to each well, and the cells were incubated at room temperature for 5 min in the dark. Fluorescence images were then acquired using a fluorescence microscope.

### Cytotoxicity Assays

Cell viability was evaluated using a Cell Counting Kit-8 (CCK-8) assay (Sangon Biotech, E606335). Cells were seeded into 96-well plates and cultured until they reached approximately 70– 80% confluence. Cells were then transfected with the indicated plasmids and subsequently treated with the corresponding anticancer agents as described above. After 72 h of drug treatment, the cells were gently washed three times with PBS. Subsequently, 100 μL of freshly prepared CCK-8 working solution, consisting of 90 μL of culture medium and 10 μL of CCK-8 Solution, was added to each well. The cells were incubated at 37°C for 30 min in the dark, and the absorbance was measured at 450 nm using a microplate reader. Cell viability was calculated according to the following formula: Cell viability (%) = [(OD (experimental) – OD (blank)]/ [ (OD (control) – OD (blank)] × 100%.

### ROS Measurement

Reactive oxygen species (ROS) levels were detected using a ROS Assay Kit (Beyotime Biotechnology, S0033S). For fluorescence microscopic analysis, the culture medium was removed, and the cells were washed three times with PBS. Subsequently, 100 μL of DCFH-DA probe at a working concentration of 10 μM was added to each well of a 96-well plate. The cells were incubated at 37°C for 20 min in the dark. After incubation, the cells were washed three times with PBS, and fluorescence images were acquired using a fluorescence microscope. For flow cytometric analysis, treated cells cultured in 6-well plates were collected and resuspended in 200 μL of DCFH-DA probe at a working concentration of 10 μM. The cells were incubated at 37°C for 20 min in the dark and gently inverted every 5 min during incubation. The cells were subsequently washed three times with PBS and analyzed using a flow cytometer. The mean fluorescence intensity (MFI) was used to quantify intracellular ROS levels.

### Iron Content Measurement

Cellular ferrous iron (Fe²⁺) levels were detected using the Cellular Red Fluorescent Ferrous Ion Assay Kit with RhoNox-6 (Beyotime Biotechnology, S1070S). For fluorescence microscopic analysis, the culture medium was removed, and the cells were washed three times with PBS. Subsequently, 100 μL of RhoNox-6 staining solution was added to each well. The cells were incubated at 37°C for 30 min in the dark, and fluorescence images were acquired using a fluorescence microscope. For flow cytometric analysis, treated cells cultured in 6-well plates were collected and resuspended in 200 μL of RhoNox-6 staining solution. The cells were incubated at 37°C for 30 min in the dark and subsequently analyzed using a flow cytometer.

### Assessment of lipid peroxidation

Lipid peroxidation was assessed using the Lipid Peroxidation Assay Kit with BODIPY™ 581/591 C11 (Beyotime Biotechnology, S0043S). For fluorescence microscopic analysis, the culture medium was removed, and the cells were washed once with PBS. Subsequently, 100 μL of BODIPY™ 581/591 C11 working solution was added to each well, followed by incubation at 37°C for 15 min. The staining solution was then removed, and the cells were washed twice with PBS before fluorescence images were acquired using a fluorescence microscope. For flow cytometric analysis, treated cells cultured in 6-well plates were collected and resuspended in 200 μL of BODIPY™ 581/591 C11 staining solution. After incubation at 37°C for 15 min, the cells were collected by centrifugation at 1,500 rpm and washed twice with PBS before analysis using a flow cytometer. Lipid peroxidation was quantified as the ratio of mean fluorescence intensity (MFI) detected in the FITC channel to that detected in the PE channel, expressed as the 530/572 nm ratio [MFI_FITC_/MFI_PE_].

### Animal treatments

Six-week-old female NOD/ShiLtJGpt-Prkdc^em26Cd52^Il2rg^em26Cd^^22^/Gpt (NCG) mice were purchased from GemPharmatech (Nanjing, China), and five-week-old female BALB/cNj-Foxn1^nu^/Gpt (BALB/c-Nude) mice were purchased from Cavens (Changzhou, China). All animal experiments were conducted in accordance with the guidelines and ethical standards of the Institutional Animal Care and Use Committee of Southeast University (Nanjing, China). Tumor growth was monitored by measuring tumor dimensions using a vernier caliper. Tumor volume was calculated using the following formula: V = (a × b²)/2, where a represents the longest tumor diameter and b represents the shortest tumor diameter. Two xenograft models were established in this study, as described below.

A human breast cancer xenograft model was established by subcutaneously inoculating MCF-7/ADR cells into the right flank of 5-week-old female BALB/c nude mice. MCF-7/ADR cells were resuspended in PBS at a concentration of 2 × 10⁸ cells/mL and mixed thoroughly with an equal volume of Matrigel (Corning, 356234). A total of 100 μL of the cell–Matrigel suspension containing 1 × 10⁷ cells was subcutaneously injected into the right flank of BALB/c nude mice. When the tumors reached approximately 150 mm³, the mice were randomly assigned to four groups: rAAV-NT, rAAV-NT + DOX, rAAV-T7, and rAAV-T7 + DOX. rAAV-NT or rAAV-T7 was administered intravenously at a dose of 1.5 × 10¹¹ vg per mouse. DOX was administered intraperitoneally at 2 mg/kg once every 3 days. Tumor volume and body weight were monitored throughout the treatment period. When the tumor volume reached approximately 800 mm³, the mice were euthanized. The heart, liver, spleen, lung, kidney, and tumor tissues were collected for subsequent H&E staining, viral DNA quantification, and miRNA expression analysis. Serum samples were collected from mice in each group for serum biochemical analysis.

A human triple-negative breast cancer xenograft model was established by subcutaneously injecting MDA-MB-231 cells into the right flank of 6-week-old female NCG mice. MDA-MB-231 cells were resuspended in PBS at a concentration of 6 × 10^7^ cells/mL and mixed thoroughly with an equal volume of Matrigel. A total of 100 μL of the resulting cell–Matrigel suspension was subcutaneously injected into the right flank of each mouse. When the tumors reached approximately 150 mm³, the mice were randomly assigned to four groups: rAAV-NT, rAAV-NT + DOX, rAAV-T7, and rAAV-T7 + DOX. rAAV-NT or rAAV-T7 was administered intravenously at a dose of 1.5 × 10¹¹ vg per mouse. DOX was administered intraperitoneally at 1 mg/kg once every 3 days. Tumor volume and body weight were monitored throughout the treatment period. When the tumor volume reached approximately 2,000 mm³, the mice were euthanized. The heart, liver, spleen, lung, kidney, and tumor tissues were collected for subsequent H&E staining, viral DNA quantification, and miRNA expression analysis. Serum samples were collected from mice in each group for serum biochemical analysis.

### Measurement of MDA levels

Malondialdehyde (MDA) levels in tissue samples were measured using an MDA Assay Reagent Kit (KeyGEN BioTECH, KGA7101) according to the manufacturer’s instructions. Briefly, 100 mg of fresh tissue was homogenized in 1 mL of pre-cooled Reagent A to prepare a 10% tissue homogenate. The homogenate was centrifuged at 12,000 × g for 5 min, and the supernatant was collected for subsequent analysis. An aliquot of 0.2 mL of the tissue homogenate supernatant was mixed with 0.2 mL of Reagent B, followed by the addition of 1.5 mL each of Reagent C and Reagent D. After thorough mixing, 1 mL of double-distilled water was added to each tube, and the samples were immediately mixed. The reaction mixtures were incubated in a boiling water bath for 50 min and rapidly cooled in an ice-water bath. The samples were then centrifuged at 3,000 rpm for 15 min at 4°C. The absorbance of the resulting supernatants was measured at 532 nm using a microplate reader.

### Measurement of tissue Fe²⁺ levels

Tissue ferrous iron (Fe²⁺) levels were measured using a Ferrous Iron Colorimetric Assay Kit (Elabscience, E-BC-K773-M). Briefly, 0.1 g of fresh tissue was homogenized in 0.9 mL of Reagent 5 and centrifuged at 12,000 × g for 10 min. The supernatant was collected for subsequent analysis. An aliquot of 100 μL of each tissue homogenate supernatant was used for protein concentration determination. For Fe²⁺ quantification, 300 μL of standard solutions at different concentrations were added to the corresponding 1.5-mL tubes to generate a standard curve. For sample measurements, 300 μL of each tissue sample was transferred into the corresponding 1.5-mL tubes. Subsequently, 150 μL of Reagent 2 was added to each tube and mixed thoroughly. The samples were incubated at 37°C for 10 min and then centrifuged at 12,000 × g for 10 min. The resulting supernatants were transferred to the corresponding wells of a microplate, and the absorbance was measured at 593 nm using a microplate reader. The Fe²⁺ content was calculated according to the standard curve and normalized to the protein concentration using the following equation: Fe²⁺ content (nmol/mg protein) = (ΔA − b) / a × f / c. ΔA=OD_sample_-OD_blank_; a: the slope of the standard curve; b: the intercept of the standard curve; f: the dilution factor of the sample before addition to the assay system; c: the protein concentration of the sample (mg/mL).

### Assessment of GSH levels in tissues

Glutathione (GSH) levels in tissue samples were measured using a Glutathione Assay Kit (KeyGEN BioTECH, KGA7305) according to the manufacturer’s instructions. Briefly, fresh tissue samples were accurately weighed and homogenized in physiological saline at a ratio of 1 g tissue to 9 mL saline under ice-cold conditions to prepare a 10% tissue homogenate. The homogenates were centrifuged at 2,500 rpm, and the resulting supernatants were collected for subsequent analysis. An aliquot of 0.5 mL of the 10% tissue homogenate supernatant was mixed thoroughly with 2 mL of Reagent 1 working solution and centrifuged at 3,500–4,000 rpm for 10 min. Subsequently, 1 mL of the resulting supernatant was collected for the colorimetric reaction. After incubation at room temperature for 5 min, the absorbance was measured at 420 nm. GSH content was calculated using the following formula: [OD_test_ -OD_blank_] / [OD_standard_ -OD_blank_] × 20 μmol/L × 307 ÷c (prot)

### Hematoxylin and eosin (H&E) staining

Freshly collected tissues, including the heart, liver, spleen, lung, kidney, and tumor tissues, were fixed in 4% paraformaldehyde (Servicebio, G1101) at room temperature for 24 h. The fixed tissues were routinely dehydrated through a graded ethanol series, cleared in xylene, and embedded in paraffin. Paraffin-embedded tissues were sectioned at a thickness of 4 μm and mounted onto glass slides. The sections were deparaffinized in xylene and rehydrated through a graded ethanol series, followed by staining with hematoxylin and eosin (H&E) (Beyotime Biotechnology, C0105S). After dehydration and clearing, the sections were mounted with neutral resin and examined under a light microscope. Representative images were captured for histopathological evaluation.

### Statistical analysis

All data are presented as the mean ± standard deviation (SD). Statistical analyses and graphing were performed using GraphPad Prism 10.0. Statistical differences between two groups were assessed using a two-tailed Student’s t-test. For comparisons involving three or more groups, one-way or two-way analysis of variance (ANOVA) was performed, followed by Tukey’s or Sidak’s multiple-comparison test, as appropriate. A P value < 0.05 was considered statistically significant.

## Supplementary Information

The online version contains supplementary material available on website. Supplementary Material 1

## Author contributions

J.W. got funding for this study and concepted and supervised this study. Y.Z. under direction of M.C. and J.W. designed and performed main experiments, analyzed the data, and wrote draft of manuscript. Y.Z., M.C., and J.W. wrote the manuscript. L.W. and F.Z. prepared PDOs. Z.K., A.Y., X.W., and X.X. performed part of data analysis and experiments. Y.S. given helpful discussion and suggestion about the selection of tumor models and chemicals from the perspective of clinical application. All authors contributed to the critical revision of the manuscript and approved the final version.

## Data availability

The authors declare that the main data supporting the findings of this study are available within the Article and the Supplementary Information or available from the corresponding author upon reasonable request. The source data underlying Figs. 1b, c, d, e, 2a, b, 3c, 4b, 5c, d, 6b, c, f, and 7b, f, d, h, i, j and Supplementary Figs. 1a, b, c, 2, 3a, 4c, 5b, 6c, 8b, 9b, 10a, b, c, 11, 12a, b, c, 13, 14, 15a, b, 16c, and 17b, e is provided with the paper as a Source Data file.

## Declarations

### Ethics approval and consent to participate

All animal experiments were performed under protocols approved by the Animal Experimentation and Ethics Committee of Southeast University. All tissues were obtained from patients who had undergone surgery at the Nanjing Hospital of Chinese Medicine Affiliated to Nanjing University of Chinese Medicine, with the consent of the patients and approval of the Nanjing Hospital of Chinese Medicine Committee for Ethical Review of Research involving Human Subjects (approval number: KY2025052). All the patients signed an informed consent form.

### Consent for publication

Not applicable.

