## Supplementary Table 1-2 and Supplementary Figures 1-17 for "Sensitize drug-resistant tumors to chemo- and targeted therapies by smart ferroptosis"

**Table S1.** Oligonucleotides sequences used to construct vector pAAV-T7.

| | |
|---|---|
| miFSP1-F | TGCTGCAAACAAACAAATAAAGTGGAGTTTTGGCCACTGACTGACTCCACTTTTTGTTTGTTTG |
| miFSP1-R | CCTGCAAACAAACAAAAAGTGGAGTCAGTCAGTGGCCAAAACTCCACTTTATTTGTTTGTTTGC |
| miFHT1-F | TGCTGTAAGGAATCTGGAAGATAGCCGTTTTGGCCACTGACTGACGGCTATCTCAGATTCCTTA |
| miFHT1-R | CCTGTAAGGAATCTGAGATAGCCGTCAGTCAGTGGCCAAAACGGCTATCTTCCAGATTCCTTAC |
| miGPX4-F | TGCTGTTCAGTAGGCGGCAAAGGCGGGTTTTGGCCACTGACTGACCCGCCTTTCGCCTACTGAA |
| miGPX4-R | CCTGTTCAGTAGGCGAAAGGCGGGTCAGTCAGTGGCCAAAACCCGCCTTTGCCGCCTACTGAAC |
| miNRF2-F | TGCTGTAAAGTAGCAGGTGAGGGCATGTTTTGGCCACTGACTGACATGCCCTCCTGCTACTTTA |
| miNRF2-R | CCTGTAAAGTAGCAGGAGGGCATGTCAGTCAGTGGCCAAAACATGCCCTCACCTGCTACTTTAC |
| miSLC7A11-F | TGCTGATAACCTGGAGACAGCAAACAGTTTTGGCCACTGACTGACTGTTTGCTCTCCAGGTTAT |
| miSLC7A11-R | CCTGATAACCTGGAGAGCAAACAGTCAGTCAGTGGCCAAAACTGTTTGCTGTCTCCAGGTTATC |
| miFPN-F | TGCTGTCTACCTGCAGCTTACATGATGTTTTGGCCACTGACTGACATCATGTACTGCAGGTAGA |
| miFPN-R | CCTGTCTACCTGCAGTACATGATGTCAGTCAGTGGCCAAAACATCATGTAAGCTGCAGGTAGAC |
| miLCN2-F | TGCTGTAATGTTGCCCAGCGTGAACTGTTTTGGCCACTGACTGACAGTTCACGGGGCAACATTA |
| miLCN2-R | CCTGTAATGTTGCCCCGTGAACTGTCAGTCAGTGGCCAAAACAGTTCACGCTGGGCAACATTAC |
| miNT-F | TGCTGAAATGTACTGCGCGTGGAGACGTTTTGGCCACTGACTGACGTCTCCACGCAGTACATTT |
| miNT-R | CCTGAAATGTACTGCGTGGAGACGTCAGTCAGTGGCCAAAACGTCTCCACGCGCAGTACATTTC |

**Table S2.** Primer sequences used for qPCR.

| Froward primers | Sequence (5'-3') | Reverse primers | Sequence (5'-3') |
|---|---|---|---|
| AAV-F | TGCATGACCAGGCTCAGCTA | AAV-R | GACAGGGAAGGGAGCAGTG |
| Mus-gGAPDH-F | GGCCACGCTAATCTCATTTT | Mus-gGAPDH-R | AAGGCGGAGTTACCAGAGGT |
| HSA-gGAPDH-F | CTGACTCAGCCCTGCAAAG | HSA-gGAPDH-R | CCTGCCTTCCTCACCTGAT |
| HSA-GAPDH-F | ATTTGGTCGTATTGGGCG | HSA-GAPDH-R | CTCGCTCCTGGAAGATGG |
| Mus-GAPDH-F | TCACCACCATGGAGAAGGC | Mus-GAPDH-R | GCTAAGCAGTTGGTGGTGCA |
| HSA-FSP1-F | CAGGACACGCAGGAGGT | HSA-FSP1-R | CTTGATGCCGGTGCAGAGAA |
| Mus-FSP1-F | CTGCCTACCGCAGTGCATT | Mus-FSP1-R | ACGCCATCATTTCTGCCCA |
| HSA-FTH1-F | CCCCCATTTGTGTGACTTCAT | HSA-FTH1-R | GCCCGAGGCTTAGCTTTCATT |
| Mus-FTH1-F | CAAGTGCGCCAGAACTACCA | Mus-FTH1-R | GCCACATCATCTCGGTCAAAA |
| HSA-GPX4-F | GAGGCAAGACCGAAGTAAACTAC | HSA-GPX4-R | CCGAACTGGTTACACGGGAA |
| Mus-GPX4-F | GATGGAGCCCATTCCTGAACC | Mus-GPX4-R | CCCTGTACTTATCCAGGCAGA |
| HSA-NRF2-F | TCAGCGACGGAAAGAGTATGA | HSA-NRF2-R | TAGCGTGTTTGAGTATTTGC |
| Mus-NRF2-F | TCTTGGAGTAAGTCGAGAAGTGT | Mus-NRF2-R | GTTGAAACTGAGCGAAAAAGGC |
| HSA-SLC7A11-F | GCGTGGGCATGTCTCTGAC | HSA-SLC7A11-R | GCTGGTAATGGACCAAAGACTT |
| Mus-SLC7A11-F | GGCACCGTCATCGGATCAG | Mus-SLC7A11-R | CTCCACAGGCAGACCAGAAAA |
| HSA-FPN-F | CACAACCGCCAGAGAGGATG | HSA-FPN-R | CACATCCGATCTCCCCAAGT |
| Mus-FPN-F | TGGAACTCTATGGAAACAGCCT | Mus-FPN-R | TGGCATTCTTATCCACCCAGT |
| HSA-LCN2-F | CCCGCAAAAGATGTATGCCA | HSA-LCN2-R | CTCACCACTCGGACGAGGTA |
| Mus-LCN2-F | TGGCCCTGAGTGTCATGTG | Mus-LCN2-R | CTCTTGTAGCTCATAGATGGTGC |
| HSA-RELA-F | CCTGGAGCAGGCTATCAGTC | HSA-RELA-R | ATGGGATGAGAAAGGACAGG |
| Mus-RELA-F | TGCGATTCCGCTATAAATGCG | Mus-RELA-R | ACAAGTTCATGTGGATGAGGC |
| U6-F | GGAACGATACAGAGAAGATTAGC | U6-R | TGGAACGCTTCACGAATTTGCG |
| miFSP1-F | CAAACAAACAAATAAAGTGGA | miFTH1-F | TAAGGAATCTGGAAGATAGCC |
| miGPX4-F | TTCAGTAGGCGGCAAAGGCGG | miNRF2-F | TAAAGTAGCAGGTGAGGGCAT |
| miSLC7A11-F | ATAACCTGGAGACAGCAAACA | miFPN-F | TCTACCTGCAGCTTACATGAT |
| miLCN2-F | TAATGTTGCCCAGCGTGAACT | | |

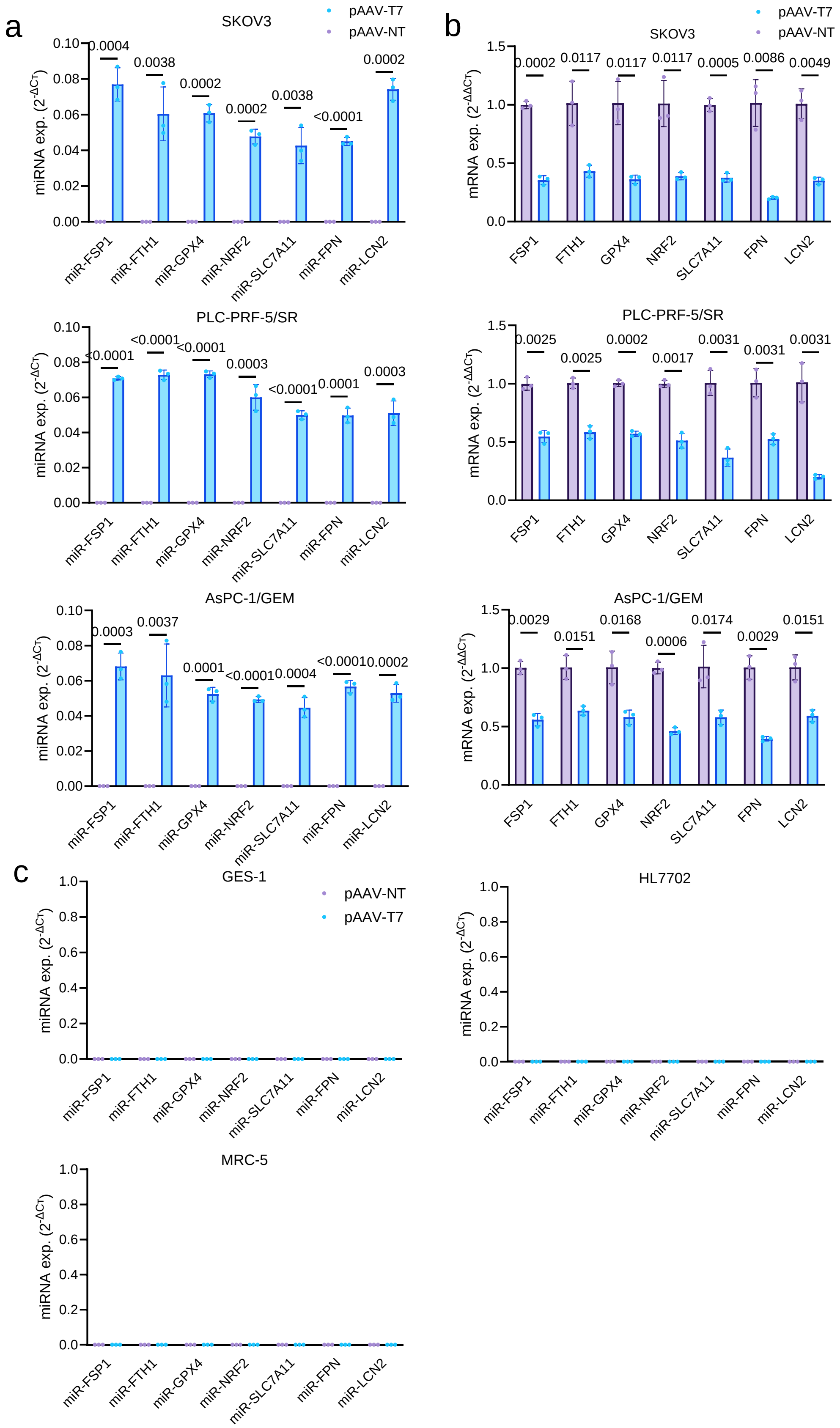


**Fig. S1 Tumor-selective expression of pAAV-T7-encoded miRNAs and suppression of their corresponding target genes.** Tumor cells (SKOV3, AsPC-1/GEM, and PLC-PRF-5/SR) and normal cells (MCF-12A, GES-1, and MRC-5) were transfected with pAAV-NT or pAAV-T7 for 48 h. (a) Expression levels of the seven pAAV-T7-encoded miRNAs, including miR-GPX4, miR-SLC7A11, miR-FSP1, miR-FPN, miR-NRF2, miR-LCN2, and miR-FTH1, in cancer cells. (b) mRNA expression levels of the corresponding target genes, including GPX4, SLC7A11, FSP1, FPN, NRF2, LCN2, and FTH1, in cancer cells. (c) Expression levels of the seven pAAV-T7-encoded miRNAs, including miR-GPX4, miR-SLC7A11, miR-FSP1, miR-FPN, miR-NRF2, miR-LCN2, and miR-FTH1, in normal cells, Gene expression was determined by qRT-PCR. Data are shown as mean ± SD (n=3wells), and $p < 0.05$ was considered statistically significant. Statistical significance was analyzed the unpaired two-tailed Student’s t-test.

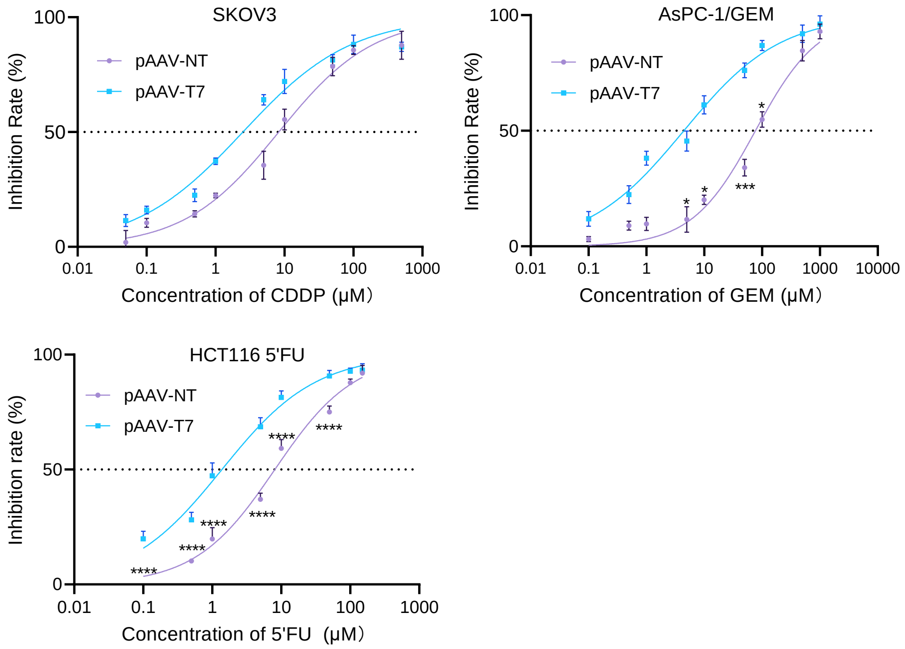


**Fig. S2. pAAV-T7 reduces the IC50 values of chemotherapeutic agents in tumor cells.** IC50 values of CDDP in SKOV3 cells, 5-FU in HCT116 cells, and GEM in AsPC-1/GEM cells following transfection with pAAV-NT or pAAV-T7. Data are presented as mean ± SD (n=3 wells). IC50 values obtained from independent experiments were calculated by nonlinear regression using a four-parameter logistic model. IC50 values between the pAAV-NT and pAAV-T7 groups were compared using a two-tailed Student's t-test. P values are given and that less than 0.05 means statistical significance.

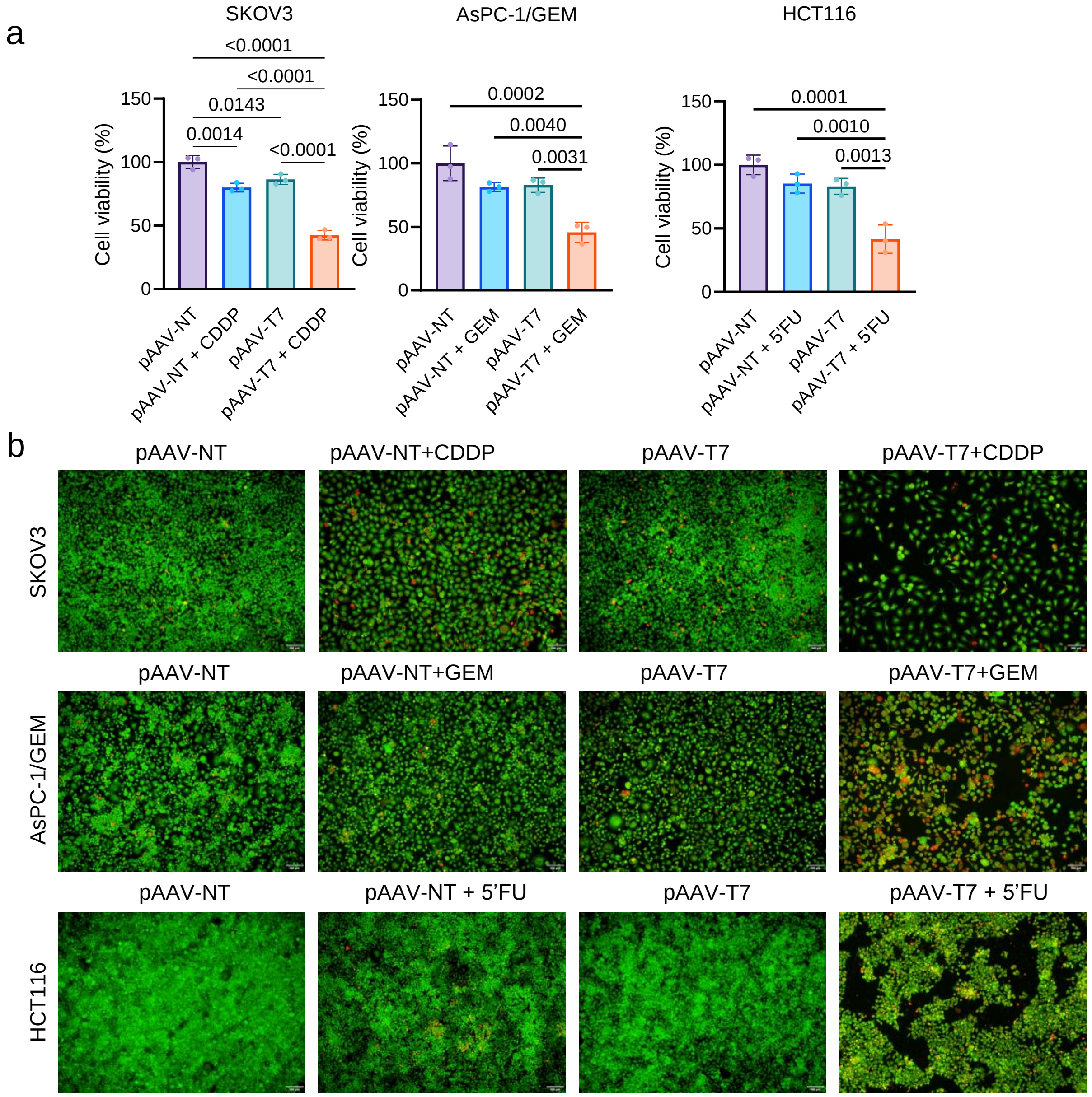


**Fig. S3. pAAV-T7 enhances chemotherapeutic agent-induced cell death and reduces cell viability.** (a) AO/EB staining of SKOV3, HCT116, and AsPC-1/GEM cells following transfection with pAAV-NT or pAAV-T7 and treatment with CDDP, 5-FU, or GEM, respectively, at fixed concentrations. (b) Cell viability assessed by CCK-8 assay under the same treatment conditions. Data are presented as mean ± SD (n=3 wells). Statistical significance was analyzed by one-way ANOVA with Tukey's test in (b). P values are given and that less than 0.05 means statistical significance.

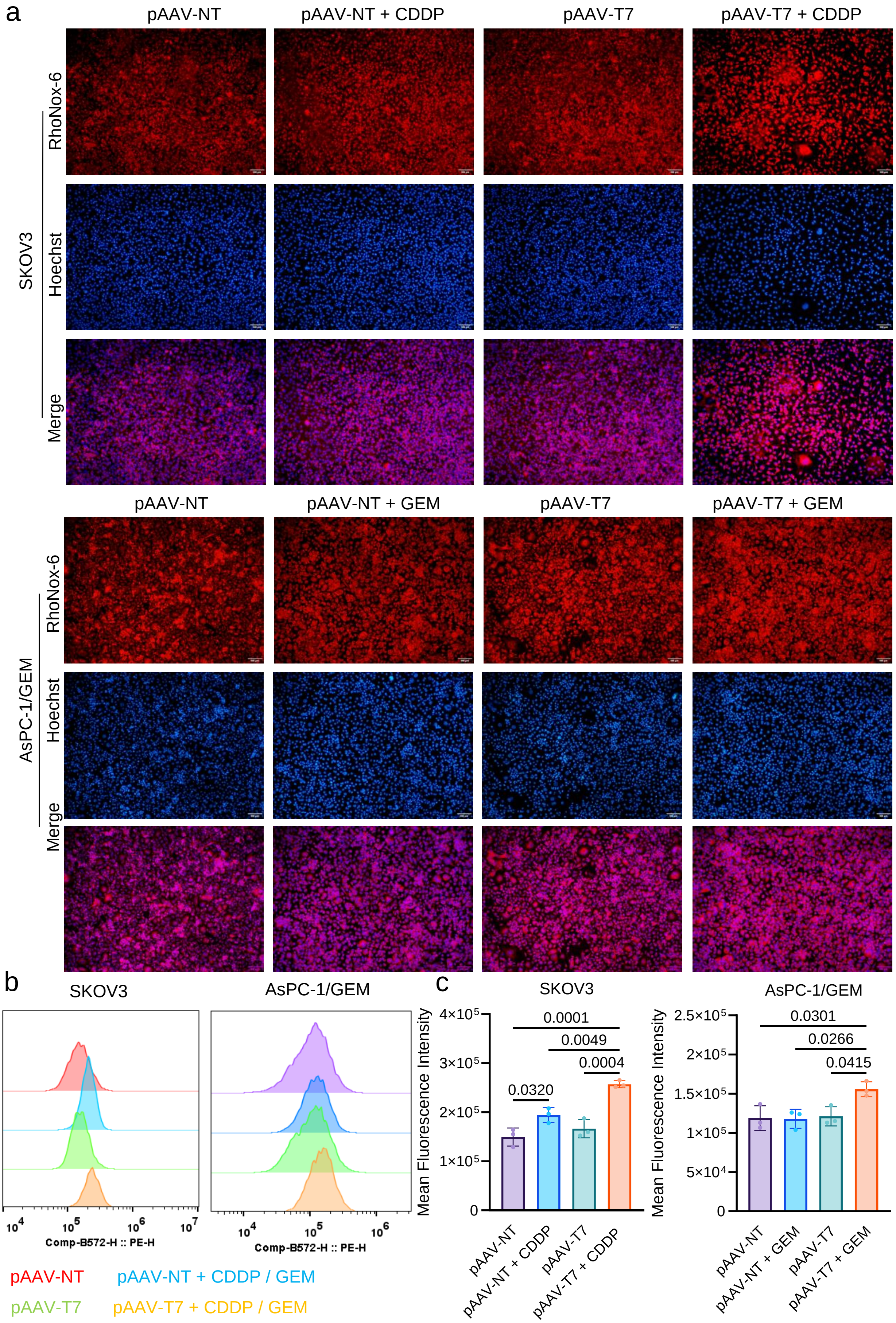


**Fig.S4 Combination of pAAV-T7 with anticancer agents promotes intracellular ferrous iron accumulation in tumor cells.** SKOV3, PLC-PRF-5/SR, and AsPC-1/GEM cells were transfected with pAAV-NT or pAAV-T7 and subsequently exposed to the corresponding anticancer agents for 48 h. (a) Representative fluorescence micrographs of intracellular $Fe^{2+}$ staining. (b) Representative flow-cytometric histograms showing intracellular $Fe^{2+}$ levels. (c) Quantification of intracellular $Fe^{2+}$ levels based on the mean fluorescence intensity (MFI) determined by flow cytometry. Data are presented as mean ± SD (n=3 independent experiments). Statistical significance was analyzed by one-way ANOVA with Tukey's test in (c). P values are given and that less than 0.05 means statistical significance.

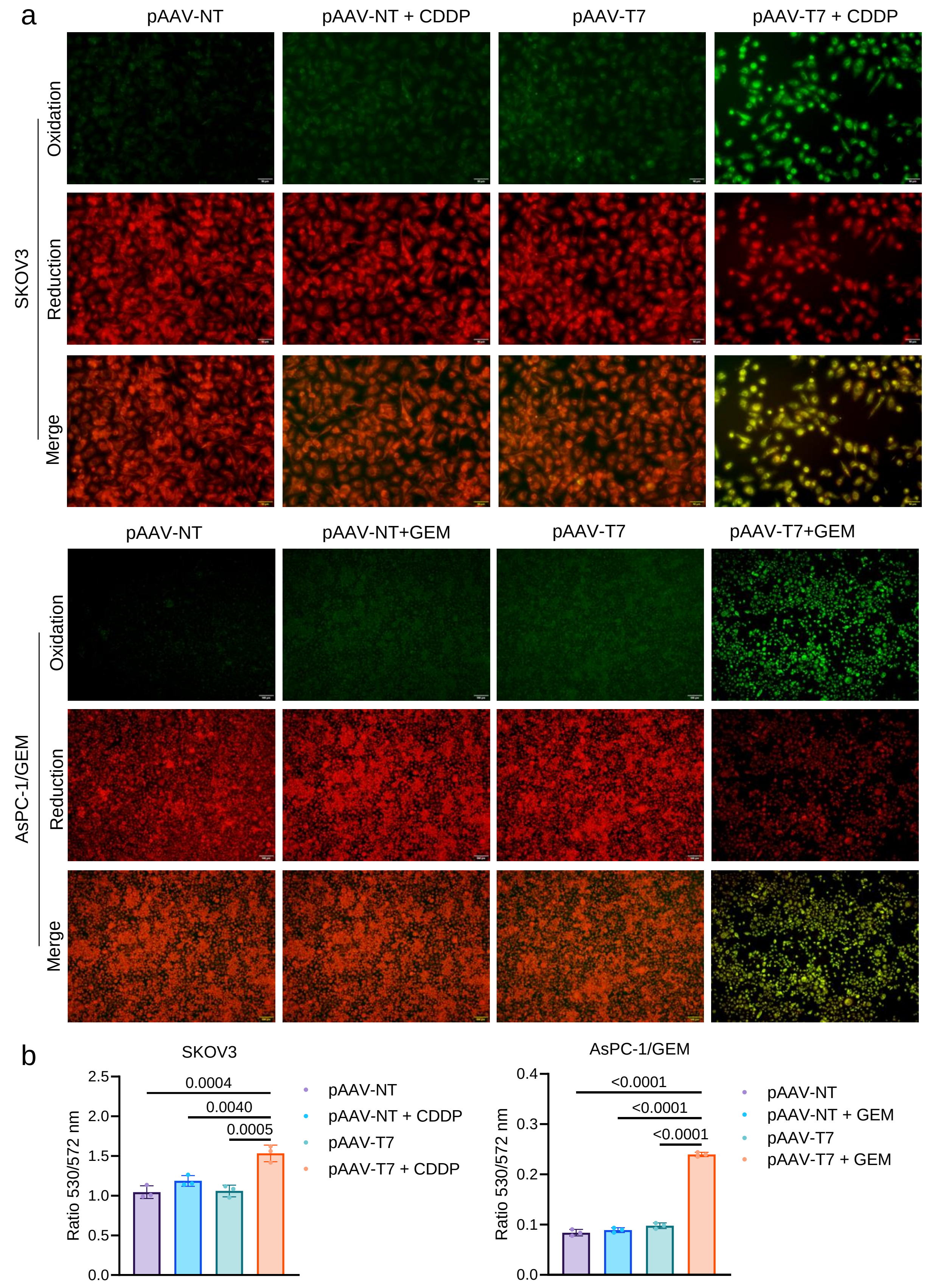


**Fig.S5 Combination of pAAV-T7 with anticancer agents promotes lipid peroxidation in tumor cells.** SKOV3, PLC-PRF-5/SR, and AsPC-1/GEM cells were transfected with pAAV-NT or pAAV-T7 and subsequently exposed to the corresponding anticancer agents for 48 h. (a) Representative fluorescence images of BODIPY 581/591 C11 staining showing lipid peroxidation in SKOV3 and AsPC-1/GEM cells. (b) Quantification of lipid peroxidation based on the fluorescence intensity of BODIPY 581/591 C11. Data are presented as mean ± SD (n=3 independent experiments). Statistical significance was analyzed by one-way ANOVA with Tukey's test in (b). P values are given and that less than 0.05 means statistical significance.

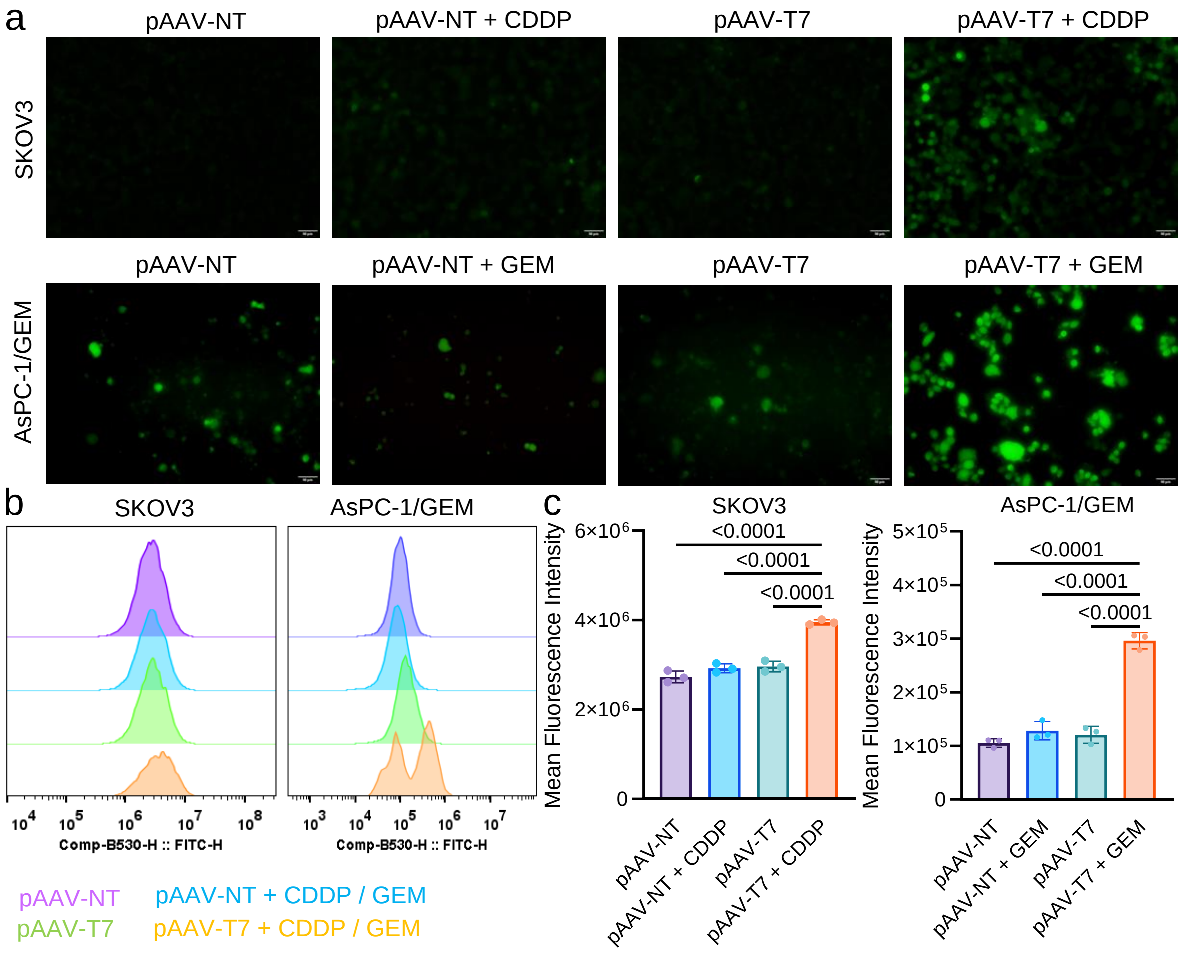


**Fig.S6 Combination of pAAV-T7 with anticancer agents increases intracellular ROS levels and induces ferroptotic cell death in tumor cells.** (a) Representative fluorescence micrographs of DCFH-DA staining showing intracellular ROS accumulation. (b) Representative flow-cytometric histograms of DCFH-DA fluorescence. (c) Quantification of intracellular ROS levels based on the mean fluorescence intensity (MFI) measured by flow cytometry. Data are presented as mean ± SD (n=3 independent experiments). Statistical significance was analyzed by one-way ANOVA with Tukey's test in (c). P values are given and that less than 0.05 means statistical significance.

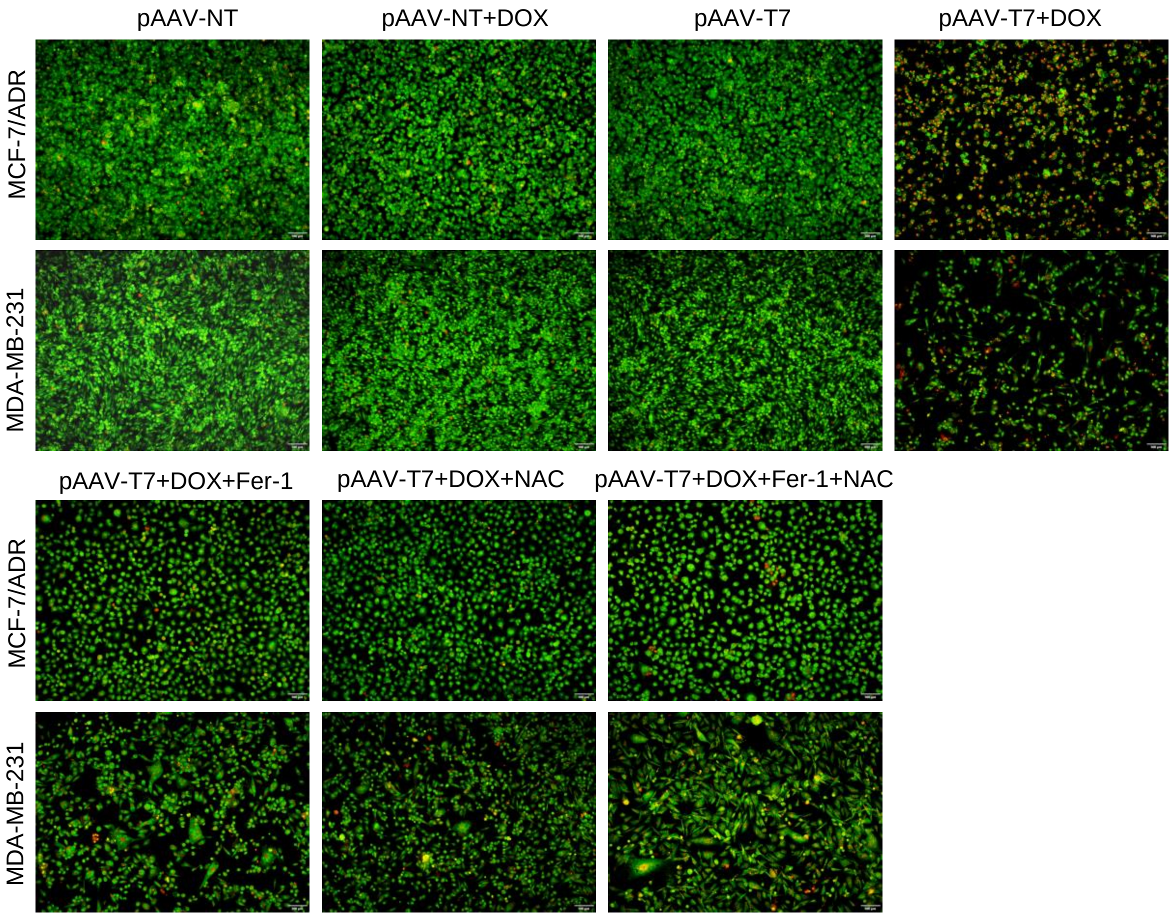


**Fig. S7. Ferrostatin-1 and NAC attenuate pAAV-T7/DOX-induced cell death in tumor cells.** AO/EB staining of MCF-7/ADR and MDA-MB-231 cells following transfection with pAAV-NT or pAAV-T7 and treatment with DOX in the presence or absence of ferrostatin-1 (Fer-1) and/or N-acetylcysteine (NAC). Representative fluorescence images of cells under the indicated treatment conditions.

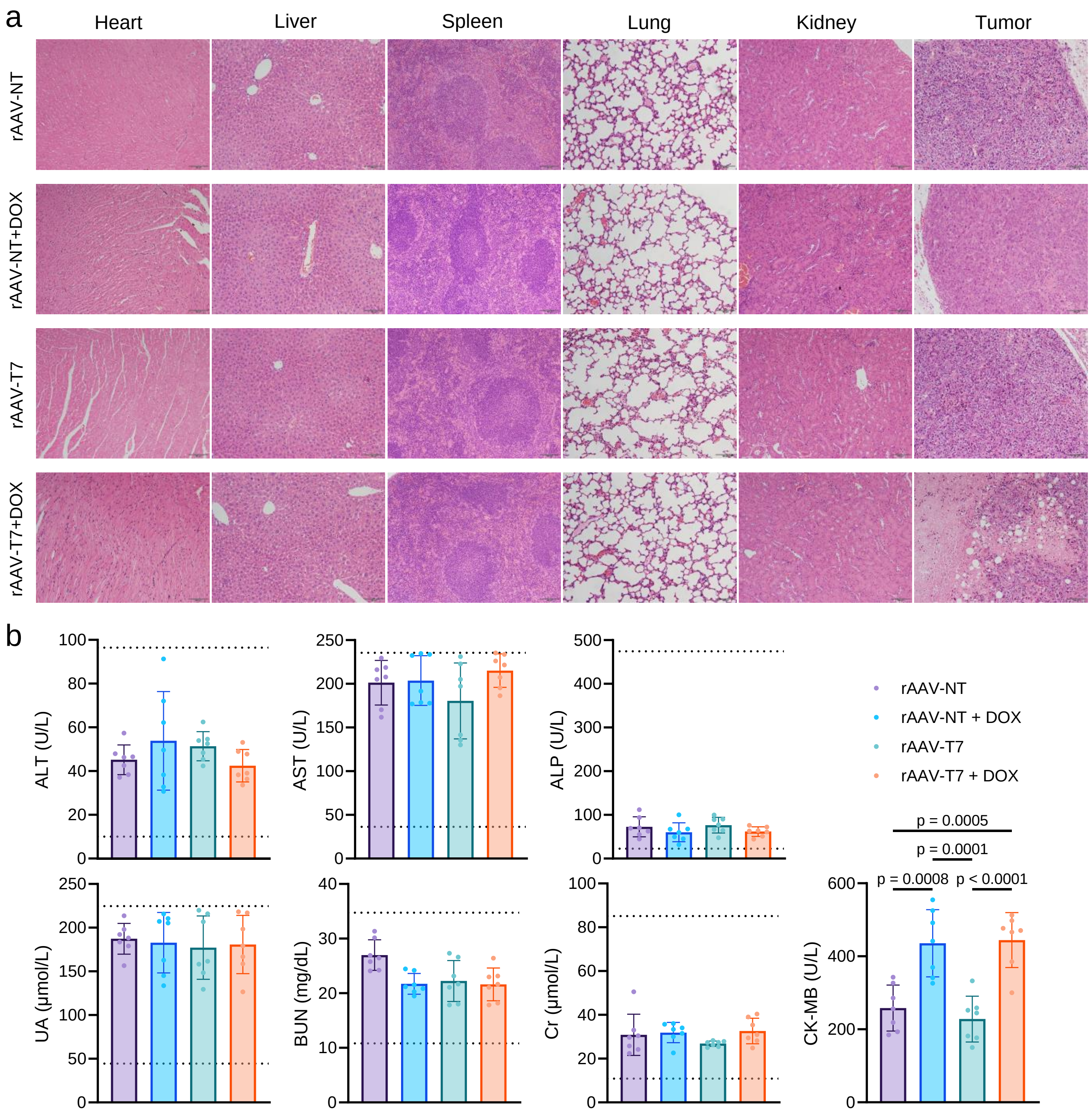


**Fig.S8 Evaluation of the in vivo safety and tumor-selective expression of rAAV-T7 in MCF-7/ADR tumor-bearing mice.** MCF-7/ADR tumor-bearing mice were treated with rAAV-NT or rAAV-T7, with or without DOX. (a) Representative hematoxylin and eosin (H&E)-stained images of the heart, liver, spleen, lung, kidney, and tumor tissues collected at the endpoint of treatment. (b) Serum biochemical parameters indicative of hepatic and renal function measured at the endpoint of treatment. Data are presented as mean ± SD (n=7). Statistical significance was analyzed by one-way ANOVA with Tukey's test. P values are given and that less than 0.05 means statistical significance.

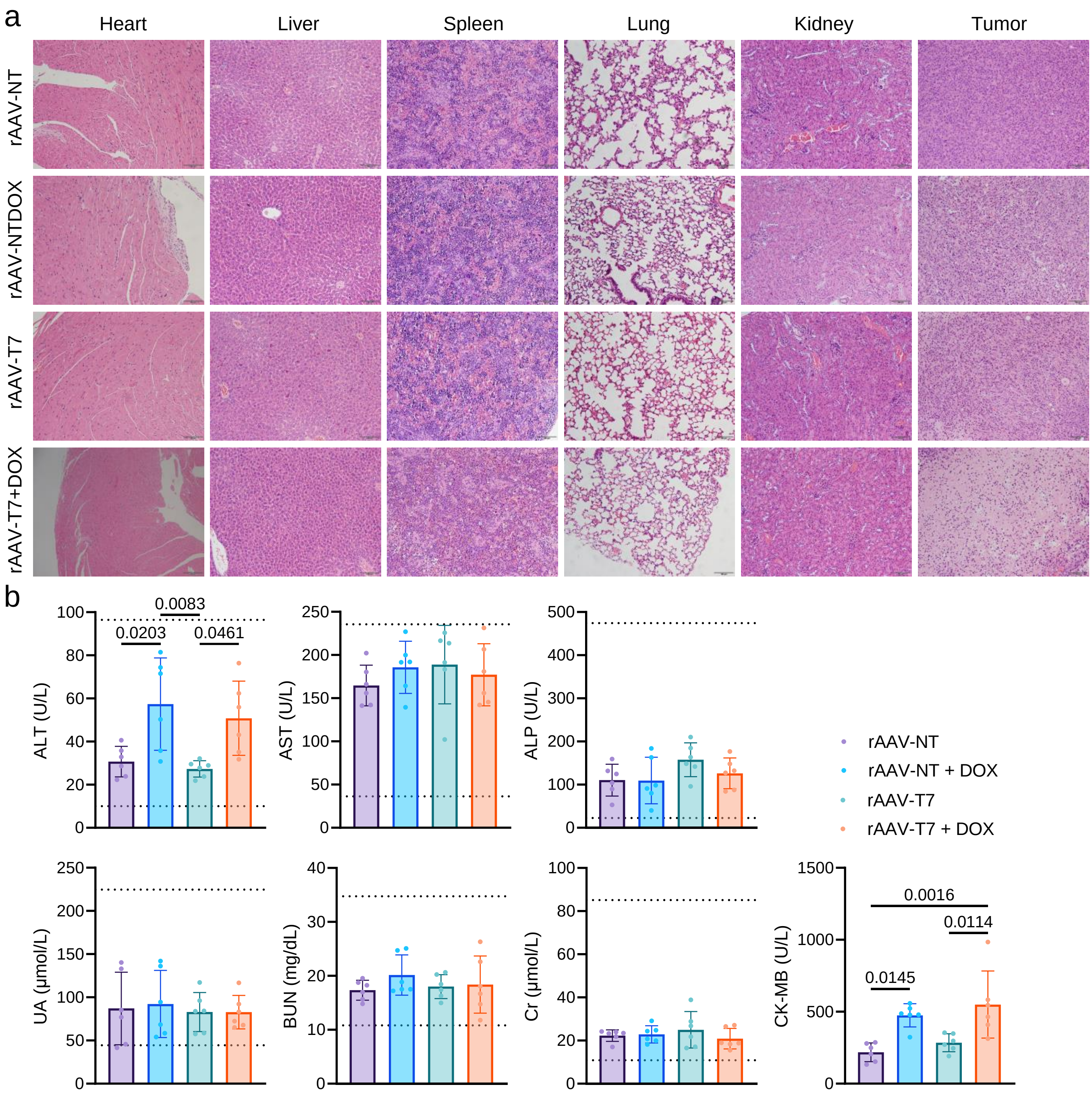


**Fig.S9 Evaluation of the in vivo safety and tumor-selective expression of rAAV-T7 in MDA-MB-231 tumor-bearing mice.** MDA-MB-231 tumor-bearing mice were treated with rAAV-NT or rAAV-T7, with or DOX. (a) Representative H&E-stained images of the heart, liver, spleen, lung, kidney, and tumor tissues collected at the endpoint of treatment. (b) Serum biochemical parameters associated with hepatic, renal, and cardiac function measured at the endpoint of treatment. Data are presented as mean ± SD (n=6). Statistical significance was analyzed by one-way ANOVA with Tukey's test. P values are given and that less than 0.05 means statistical significance.

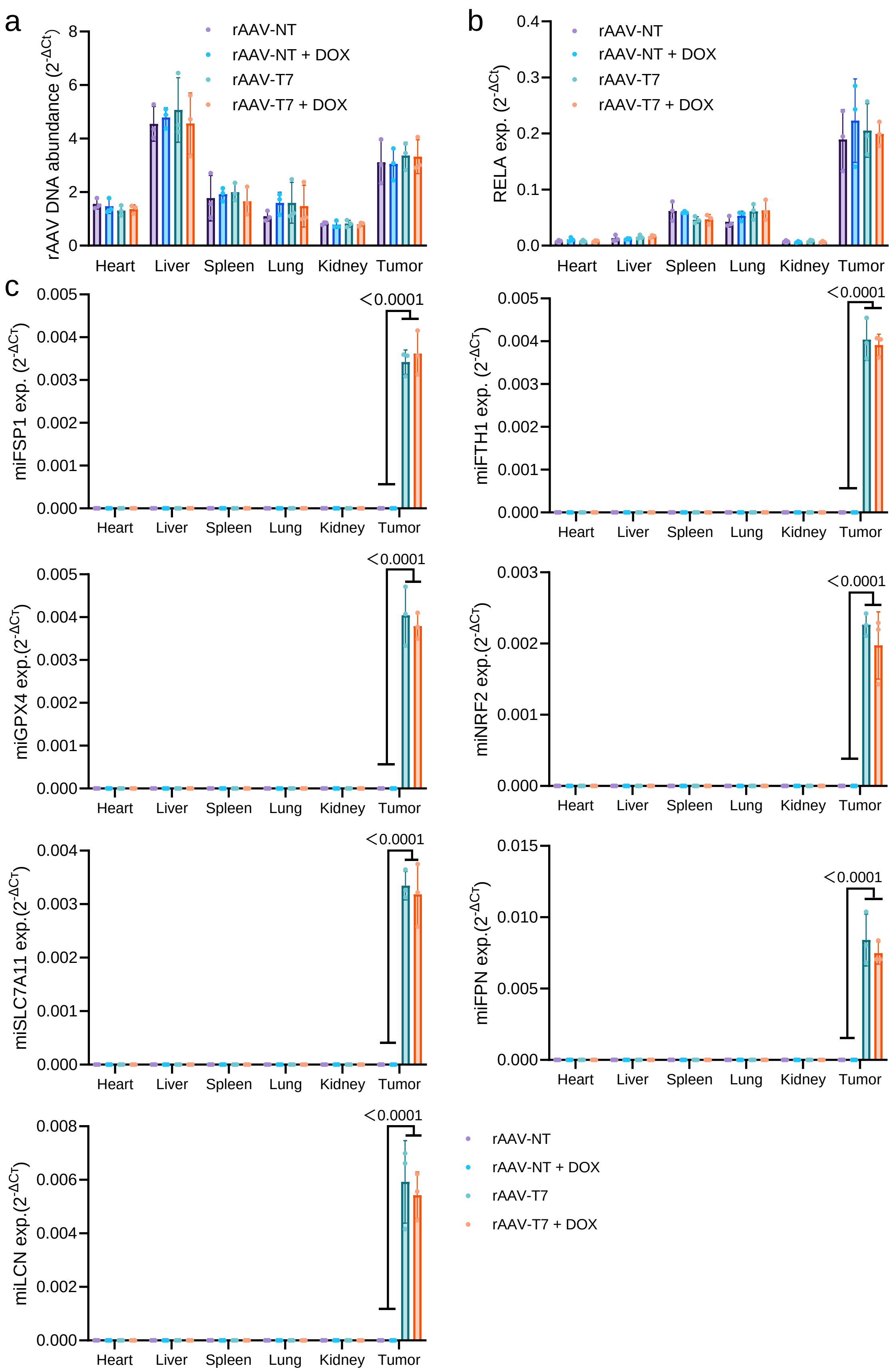


**Fig.S10 Evaluation of the in vivo safety and tumor-selective expression of rAAV-T7 in MCF-7/ADR tumor-bearing mice.** MCF-7/ADR tumor-bearing mice were treated with rAAV-NT or rAAV-T7, with or without DOX. (a) Abundance of rAAV DNA in the tissues of MCF-7/ADR tumor-bearing mice. (b) Relative mRNA expression levels of RELA in the tissues of MCF-7/ADR tumor-bearing mice. (c) Expression levels of the seven rAAV-T7-encoded miRNAs in the heart, liver, spleen, lung, kidney, and tumor tissues. Gene expression was determined by qRT-PCR. Data are presented as mean ± SD (n=7). Statistical significance was analyzed by one-way ANOVA with Tukey's test. P values are given and that less than 0.05 means statistical significance.

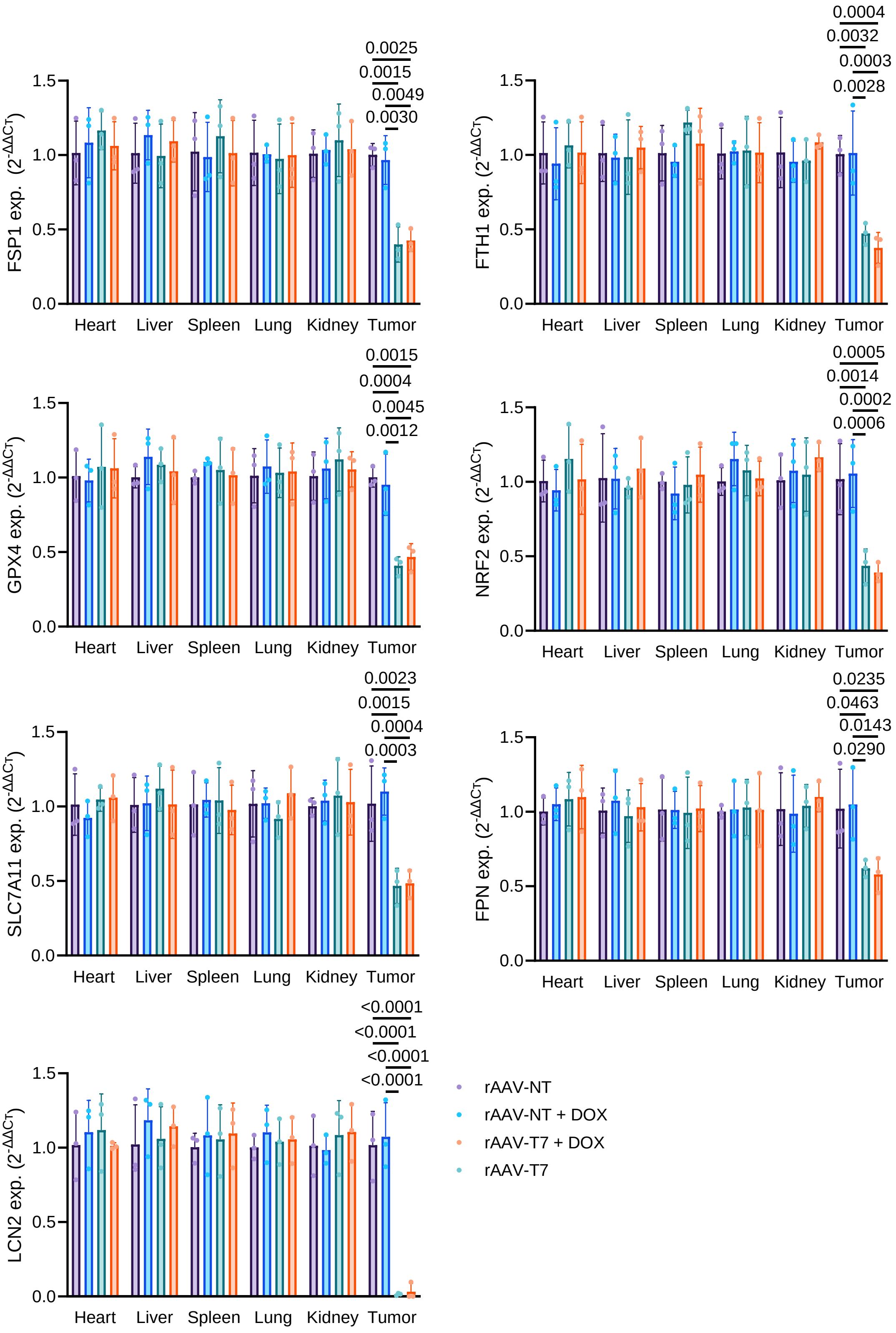


**Fig.S11 Tissue-specific expression of rAAV-T7-responsive genes and biodistribution of rAAV vectors in tumor-bearing mice.** Relative mRNA expression levels of the rAAV-T7 target genes, including GPX4, SLC7A11, FSP1, FPN, NRF2, LCN2, and FTH1, in the tissues of MCF-7/ADR tumor-bearing mice. Gene expression was determined by qRT-PCR. Data are presented as mean ± SD (n=7). Statistical significance was analyzed by one-way ANOVA with Tukey's test. P values are given and that less than 0.05 means statistical significance.

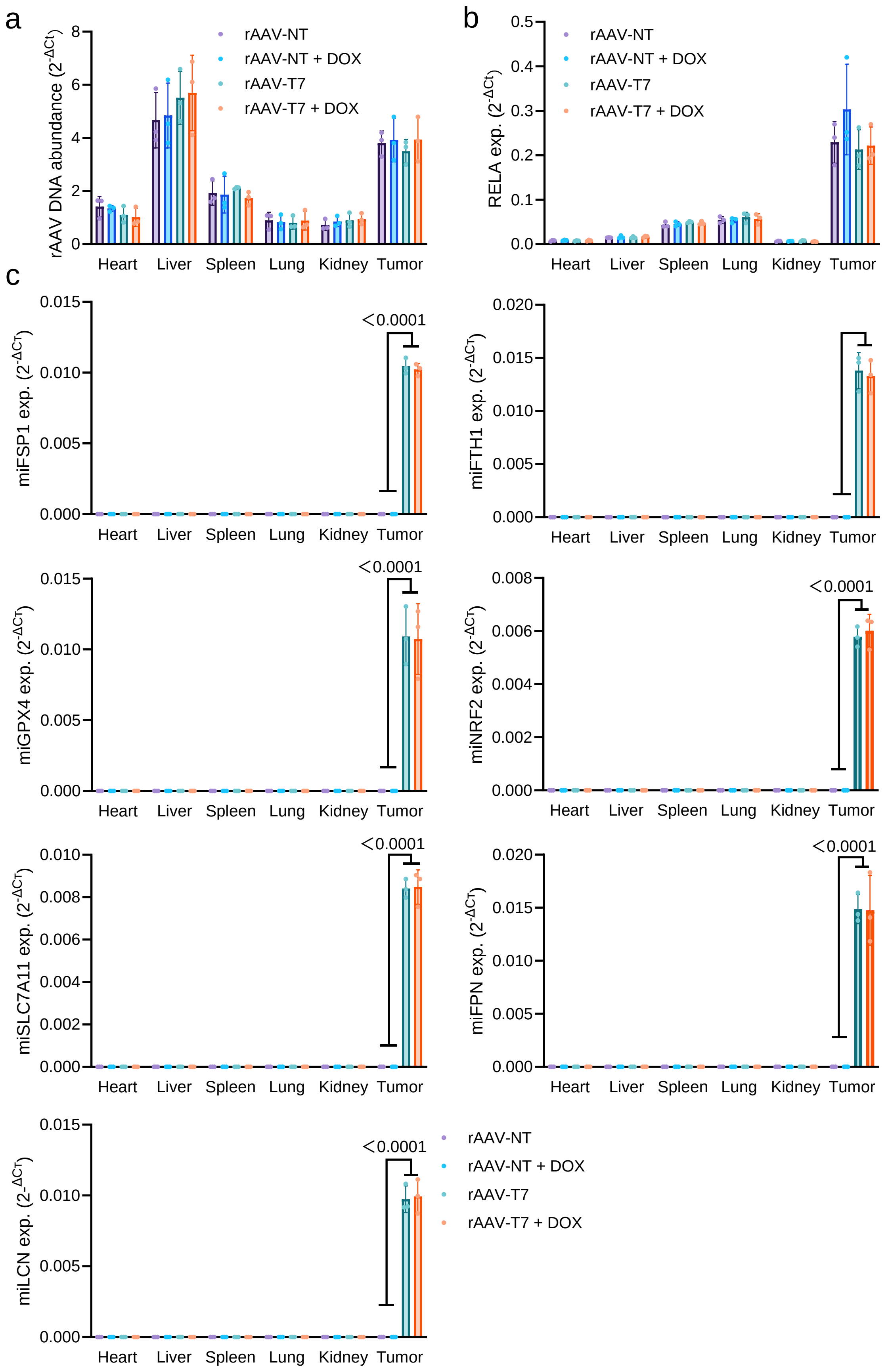


**Fig.S12 Evaluation of the in vivo safety and tumor-selective expression of rAAV-T7 in MDA-MB-231 tumor-bearing mice.** MDA-MB-231 tumor-bearing mice were treated with rAAV-NT or rAAV-T7, with or without DOX. (a) Abundance of rAAV DNA in the tissues of MDA-MB-231 tumor-bearing mice. (b) Relative mRNA expression levels of RELA in the tissues of MDA-MB-231 tumor-bearing mice. (c) Expression levels of the seven rAAV-T7-encoded miRNAs in the heart, liver, spleen, lung, kidney, and tumor tissues. Gene expression was determined by qRT-PCR. . Data are presented as mean ± SD (n=6). Statistical significance was analyzed by one-way ANOVA with Tukey’s test. P values are given and that less than 0.05 means statistical significance.

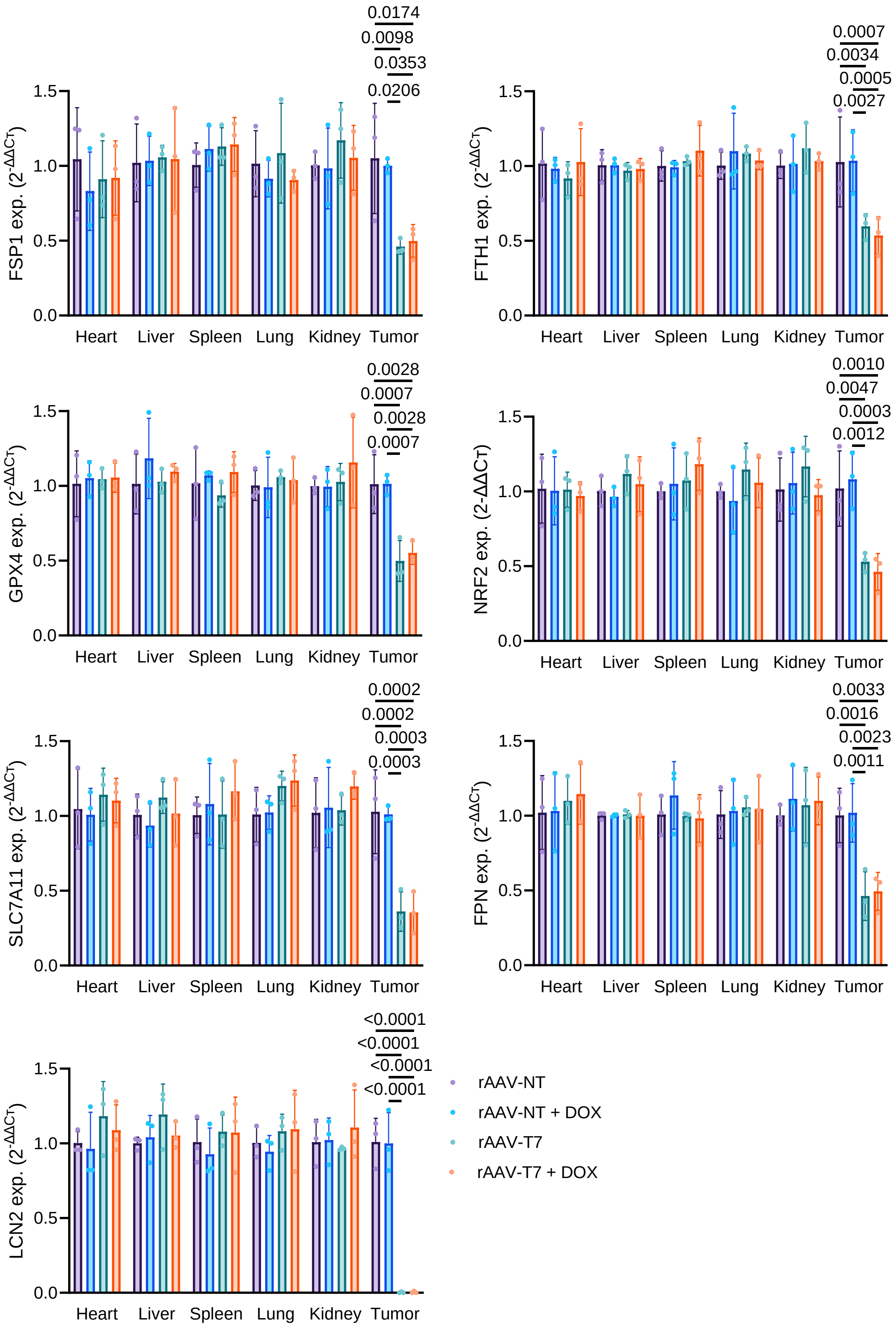


**Fig.S13 Tissue-specific expression of rAAV-T7-responsive genes and biodistribution of rAAV vectors in tumor-bearing mice.** Relative mRNA expression levels of the rAAV-T7 target genes, including GPX4, SLC7A11, FSP1, FPN, NRF2, LCN2, and FTH1, in the tissues of MDA-MB-231 tumor-bearing mice. Gene expression was determined by qRT-PCR. Data are presented as mean ± SD (n=6). Statistical significance was analyzed by one-way ANOVA with Tukey's test. P values are given and that less than 0.05 means statistical significance.

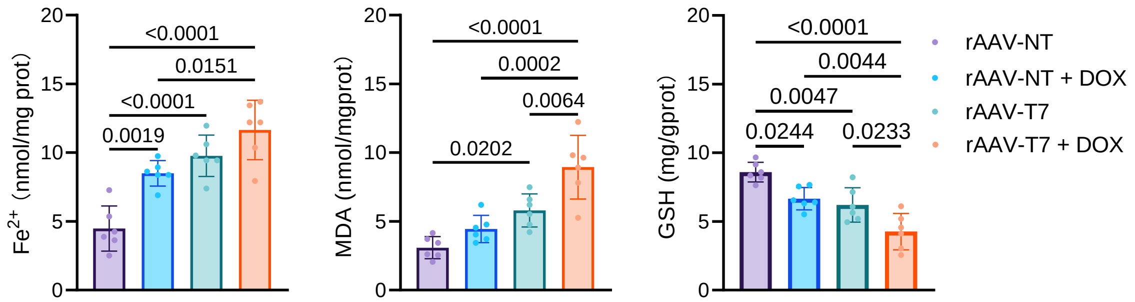


**Fig.S14 rAAV-T7 enhances ferroptosis-associated biochemical changes in MDA-MB-231 xenograft tumors.** MDA-MB-231 tumor-bearing mice were treated with rAAV-NT or rAAV-T7, with or DOX. Levels of intracellular ferrous iron ($Fe^{2+}$), malondialdehyde (MDA), and glutathione (GSH) in MDA-MB-231 xenograft tumor tissues at the endpoint of treatment. Data are presented as mean ± SD (n=6). Statistical significance was analyzed by one-way ANOVA with Tukey's test. P values are given and that less than 0.05 means statistical significance.

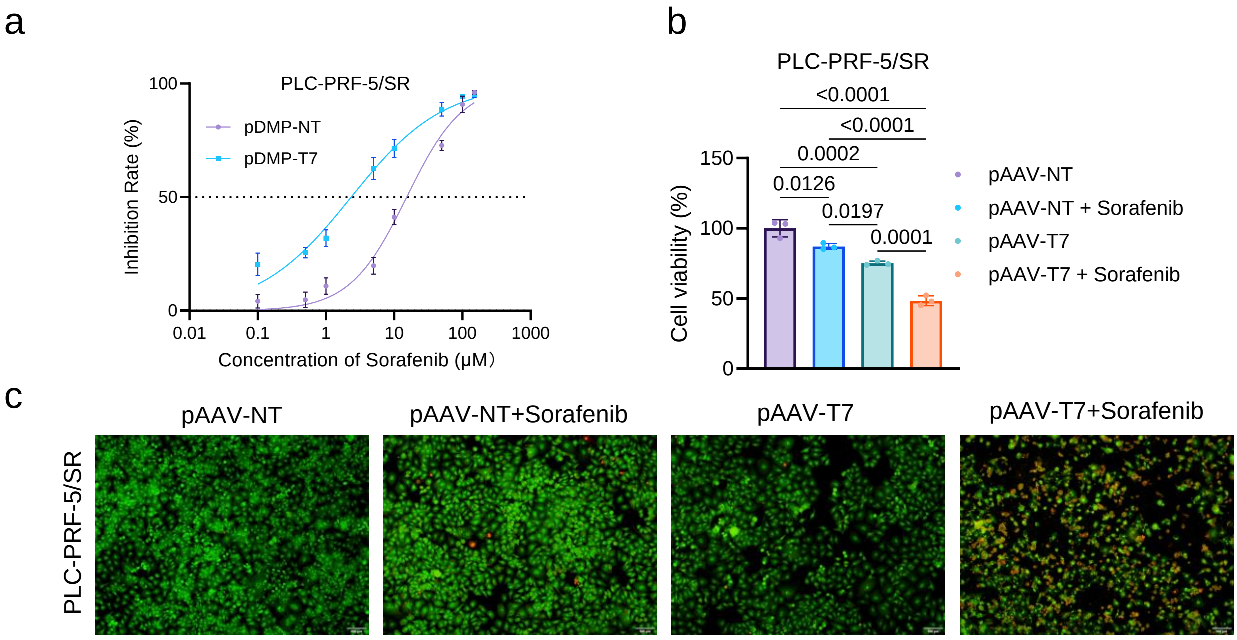


**Fig.S15 pAAV-T7 sensitizes sorafenib-resistant PLC-PRF-5/SR cells to sorafenib.** (a) IC50 values of sorafenib in PLC-PRF-5/SR cells following transfection with pAAV-NT or pAAV-T7. (b) Cell viability assessed by CCK-8 assay following treatment with sorafenib at a fixed concentration in cells transfected with pAAV-NT or pAAV-T7. (c) AO/EB staining showing cell death under the same treatment conditions. Data are presented as mean ± SD (n=3 wells). Statistical significance was analyzed by one-way ANOVA with Tukey's test in (b). IC50 values obtained from independent experiments were calculated by nonlinear regression using a four-parameter logistic model. IC50 values between the pAAV-NT and pAAV-T7 groups were compared using a two-tailed Student's t-test. P values are given and that less than 0.05 means statistical significance.

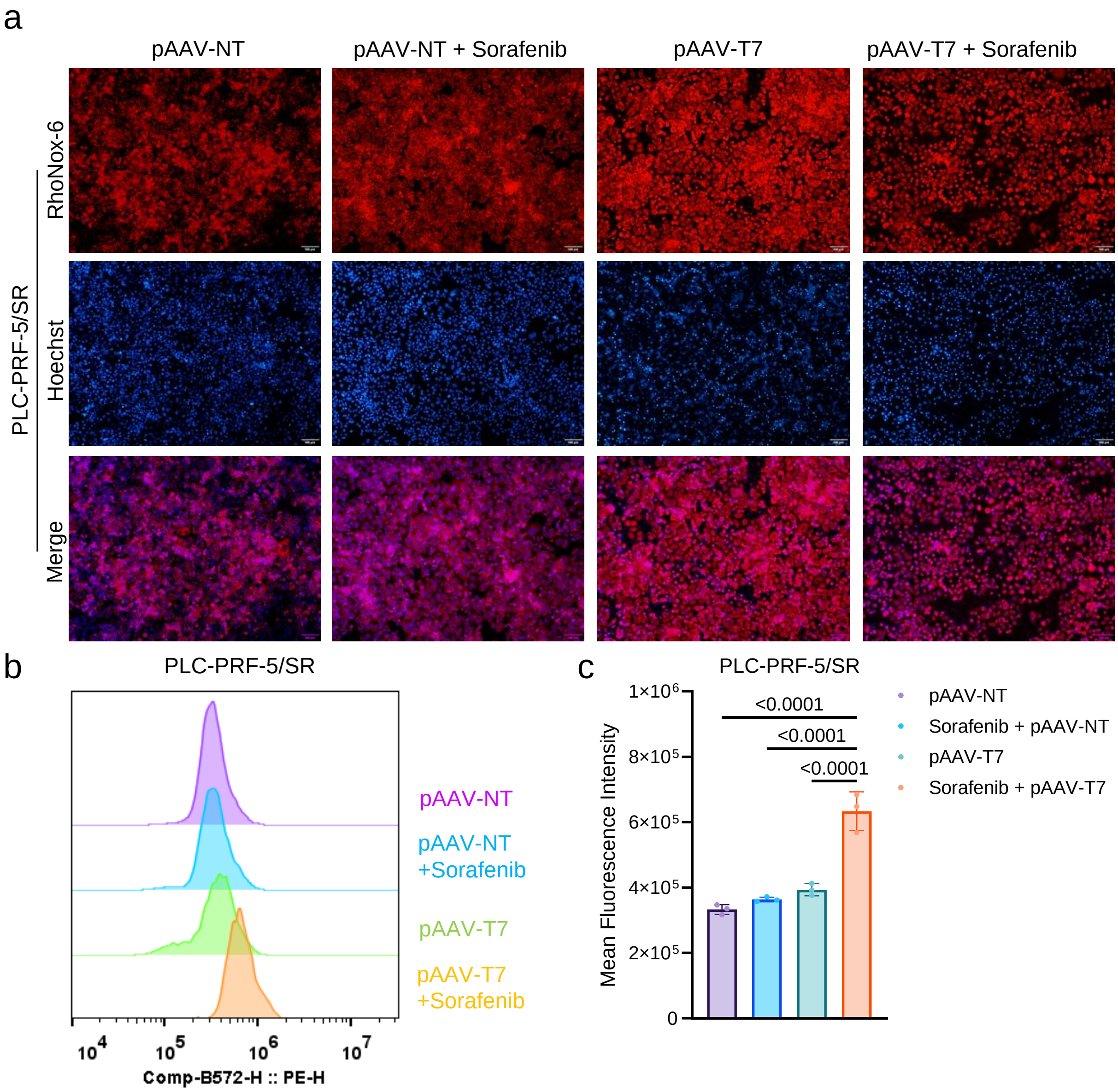


**Fig.S16 Combination treatment with pAAV-T7 and sorafenib promotes $Fe^{2+}$ accumulation in PLC-PRF-5/SR cells.** (a) Representative fluorescence images of intracellular ferrous iron ($Fe^{2+}$) in PLC-PRF-5/SR cells following transfection with pAAV-NT or pAAV-T7 and treatment with sorafenib. (b) Representative flow cytometry histograms showing intracellular $Fe^{2+}$ levels under the indicated treatment conditions. (c) Quantification of mean fluorescence intensity (MFI) of intracellular $Fe^{2+}$ detected by flow cytometry. Data are presented as mean ± SD from three independent experiments. Statistical significance was analyzed by one-way ANOVA with Tukey's test in (c). P values are given and that less than 0.05 means statistical significance.

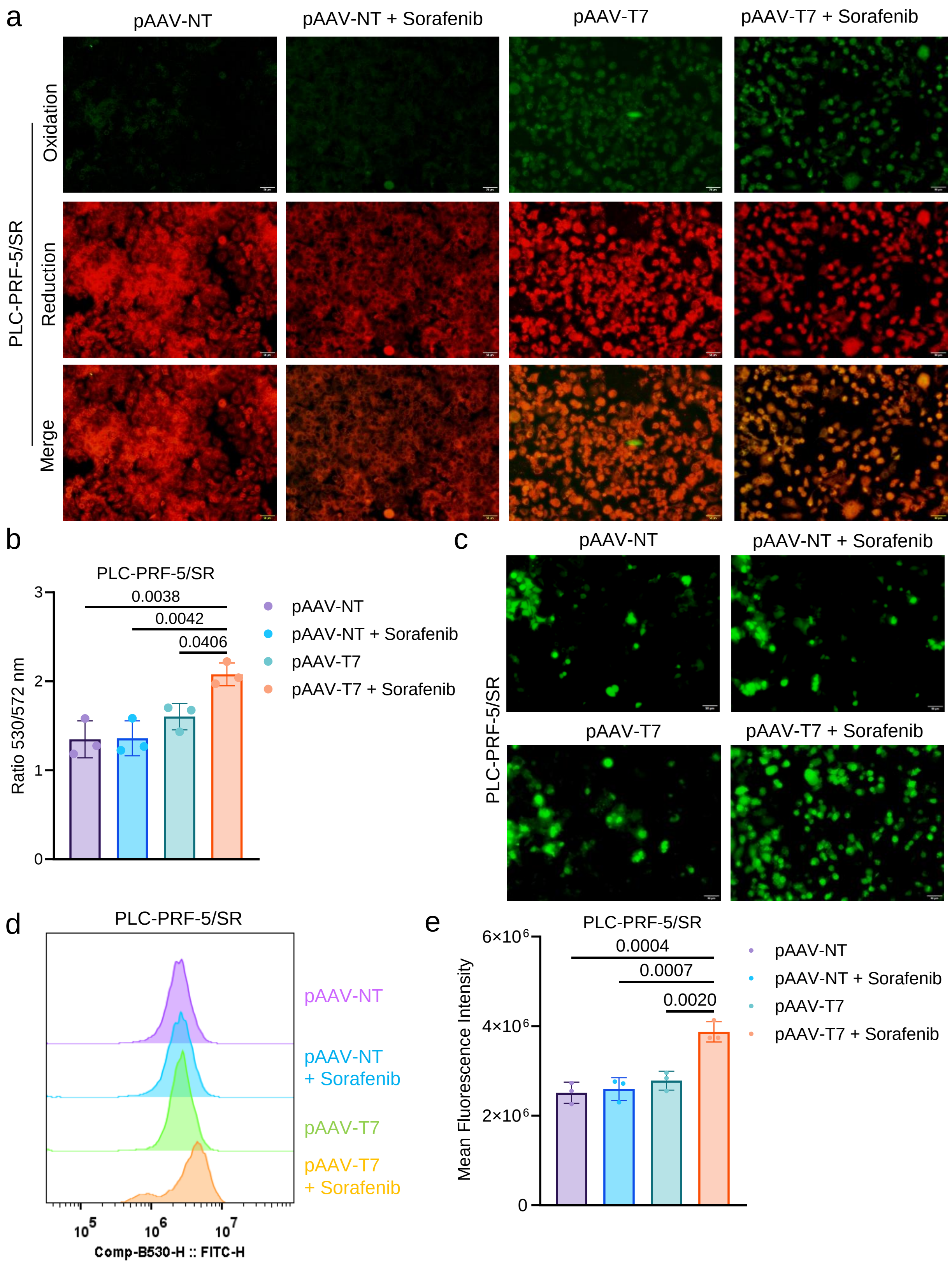


**Fig.S17 Combination treatment with pAAV-T7 and sorafenib promotes lipid peroxidation and ROS accumulation in PLC-PRF-5/SR cells.** (a) Representative fluorescence images of lipid peroxidation detected using BODIPY™ 581/591 C11 in PLC-PRF-5/SR cells following transfection with pAAV-NT or pAAV-T7 and treatment with sorafenib. (b) Quantification of lipid peroxidation based on the 530/572 nm fluorescence ratio. (c) Representative fluorescence images of intracellular total ROS detected using DCFH-DA under the indicated treatment conditions. (d) Representative flow cytometry histograms showing intracellular total ROS levels. (e) Quantification of mean fluorescence intensity (MFI) of intracellular total ROS detected by flow cytometry. Data are presented as mean ± SD from three independent experiments. Statistical significance was analyzed by one-way ANOVA with Tukey's test. P values are given and that less than 0.05 means statistical significance.
